# A new kinetic method for measuring agonist efficacy and ligand bias using high resolution biosensors and a kinetic data analysis framework

**DOI:** 10.1101/772293

**Authors:** Sam R.J. Hoare, Paul H. Tewson, Anne Marie Quinn, Thomas E. Hughes

## Abstract

The kinetics/dynamics of signaling are of increasing value for G-protein-coupled receptor therapeutic development, including spatiotemporal signaling and the kinetic context of biased agonism. Effective application of signaling kinetics to developing new therapeutics requires reliable kinetic assays and an analysis framework to extract kinetic pharmacological parameters. Here we describe a platform for measuring arrestin recruitment kinetics to GPCRs using a high quantum yield, genetically encoded fluorescent biosensor, and a new analysis framework to quantify the recruitment kinetics. The sensor enabled high temporal resolution measurement of arrestin recruitment to the angiotensin AT_1_ and vasopressin V_2_ receptors. The analysis quantified the initial rate of arrestin signaling (*k*_*τ*_), a biologically-meaningful kinetic drug efficacy parameter, by fitting time course data using routine curve-fitting methods. Biased agonism was assessed by comparing *k*_*τ*_ values for arrestin recruitment with those for Gq signaling via the AT_1_ receptor. The *k*_*τ*_ ratio values were in good agreement with bias estimates from existing methods. This platform potentially improves and simplifies assessment of biased agonism because the same assay modality is used to compare pathways (potentially in the same cells), the analysis method is parsimonious and intuitive, and kinetic context is factored into the bias measurement.

## INTRODUCTION

The kinetics / dynamics of G-protein-coupled receptor (GPCR) signaling is of increasing interest in elaborating the biology and therapeutic potential of these receptors ^1-3^. The time frame of GPCR signal transduction is dependent on the signaling pathway, regulation of signaling mechanism, and location of the receptor in the cell (spatiotemporal signaling). Temporal dynamics of signaling are being elucidated and applied to develop new therapeutics. For example, the parathyroid hormone 1 (PTH1) receptor can signal persistently over time after partitioning into the endosomal compartment ^4^. This effect was ligand dependent; PTH produced persistent signaling whereas PTH-related protein did not ^4^. This behavior is potentially involved in the therapeutic mode of action; continuous administration of PTH1 receptor agonists results in bone loss, whereas intermittent administration results in net bone formation ^5^. This kinetic effect was exploited in the development of new bone anabolic agents for treating osteoporosis ^6^. Persistent signaling is potentially beneficial for other GPCR therapeutics ^1^, for example sphingosine 1-phosphate receptor-1 agonists for treatment of multiple sclerosis ^7^.

Of potential concern, the kinetics of signaling can affect measurement of biased agonism, by affecting classical measurements of agonist activity (potency and efficacy) ^3,8^. Biased agonism is the capacity of a ligand to selectively activate one or more of multiple signaling pathways transduced by the GPCR ^9^. This concept is of considerable current interest in the development of next-generation GPCR therapeutics because it enables selective targeting towards beneficial pathways and away from potentially deleterious ones ^10-12^. For a series of dopamine receptor ligands, it was shown that the extent of bias was dependent on the time point at which the signaling responses were measured ^8^. This complicates the interpretation of bias and its translation to in vivo efficacy because it isn’t straightforward to select the most appropriate time point for establishing structure-activity relationships and for predicting in vivo efficacy from in vitro bias measurements. Differences in assay timing might contribute to discrepancies of bias estimates reported in the literature. For example, aripiprazole has been reported to be an arrestin-biased ligand but the efficacy varies from 10 to 100% and the time point from 5 minutes to 20 hours ^13-17^.

Quantifying the kinetics of signaling with useful drug parameters would aid the development of kinetically-optimized molecules, tuned, for example, to the optimum duration of signaling, timeframe of desensitization, and residence period in signaling compartments. This requires appropriately optimized kinetic assays and a data analysis platform for extracting drug parameter values from time course data. Biosensor assays have enabled high-throughput kinetic measurement of GPCR signal transduction because the same plate / well can be measured repeatedly over time ^18-20^. The signaling molecule of interest interacts with an engineered protein, changing its optical properties, for example fluorescence intensity, which is detected in specialized plate readers. Previously we and others have developed genetically-encoded biosensors incorporating fluorescent proteins such as mNeonGreen that provide high-resolution kinetic data for G-protein-mediated signals (for example, cAMP ^21^, diacylglycerol ^21^ (DAG), and Ca^2+^ ^22^). Regarding the data analysis, drug activity metrics are required which quantify the kinetics in terms that can be applied in establishing structure-activity relationships ^23^. We recently developed a data analysis framework for G-protein and downstream signaling that quantifies kinetics in terms of the initial rate of signaling ^24,25^. This rate, analogous to the initial rate of enzyme activity, is the rate of signaling before it is impacted by regulation of signaling mechanisms such as receptor desensitization and signal decay ^25^. This parameter, termed *k*_*τ*_, provides a biologically meaningful kinetic metric of ligand efficacy that has been applied to quantify ligand activity for G-protein activation and second-messenger generation ^24,25^.

The goal of this study was to optimize and integrate the biosensor modality and the data analysis framework to create a unified platform suitable for robustly measuring and quantifying signaling kinetics and biased agonism for numerous GPCR pathways. This first required extending the framework described above to arrestin signaling, since our biosensor and analysis technologies were developed only for G-protein signaling. Arrestin signaling is an alternative pathway by which GPCRs modulate cellular activity ^26,27^ and arrestin signaling has been implicated in potentially beneficial and deleterious physiological processes. For example, arrestin signaling by the angiotensin AT_1_ receptor improves the cardiac performance of ligands targeting the receptor in animal models ^28-31^, whereas arrestin signaling by the mu opioid receptor has been implicated in opioid side effects, including tolerance, reward and respiratory depression ^32-35^. This has stimulated the development of ligands biased towards or away from the arrestin pathway ^9,11,36^. Here we describe a new arrestin biosensor, utilizing mNeonGreen, suitable for generating time course data with high enough temporal resolution for the arrestin recruitment kinetics to be measured accurately. We then extended the data analysis framework to incorporate arrestin recruitment to the receptor. Finally, we applied this unified platform to directly compare arrestin and G-protein signaling via the AT_1_ receptor, using near-identical experimental conditions and the same conceptual kinetic data analysis framework. This work demonstrated a new approach for quantifying biased agonism in kinetic terms using a unified assay modality.

## RESULTS

In this study an experimental and analytical platform was developed to quantify the kinetics of arrestin recruitment in such a way as to enable direct comparison with the kinetics of downstream signaling. A new genetically-encoded biosensor that converts the change in arrestin conformation upon receptor interaction to a change in fluorescence intensity made it possible to collect detailed recordings of the kinetics of arrestin recruitment, i.e. a large number of reads at closely-spaced time points. Robust arrestin responses were obtained for the angiotensin AT_1_ and vasopressin V_2_ receptors. The time course data, i.e. the change in fluorescence intensity over time, was then analyzed using a new pharmacological analysis. This analysis quantifies the initial rate of arrestin recruitment to the receptor. The analysis was applied to responses activated by several agonists of the AT_1_ angiotensin receptor. Biased agonism was then assessed; the initial rate of arrestin recruitment was compared with the initial rate of downstream signaling measured using the same biosensor modality (diacylglycerol and Ca^2+^ mobilization). The resulting bias ratios were in good agreement with values obtained using conventional methods, validating the new method.

### Biosensor of arrestin-receptor interaction

Accurately quantifying signaling over time with an optical biosensor requires certain physical criteria. GPCR signaling is often rapid, for example the rise phase occurs within a few seconds for Ca^2+^ mobilization and within a few minutes for cAMP generation and arrestin recruitment. Consequently, in order to obtain sufficient data points, the read time, the time required to obtain sufficient signal, needs to be short, ideally < 10 seconds. Second, a large number of time points are required to accurately define the curve shape and reliably fit the relevant equations to the time course data. This requires minimal photobleaching of the sensor. These criteria can be met with direct fluorescence sensors, owing to the high quantum yield ^21,22^. This property enables short read times to be used because the signal per unit time is high. It also minimizes photobleaching because a short excitation time can be used.

We and others have developed fluorescent biosensors in which intrinsically fluorescent protein sensors have been incorporated. (Three examples used in this study are R-GECO ^22^ for Ca^2+^, Downward DAG for diacylglycerol and cADDis for cAMP ^21^.) The protein sensors have been engineered to be conformationally-sensitive, such that interaction with the signaling molecule of interest changes the optical properties, enabling the interaction to be detected as a change of fluorescence intensity. Since the proteins are intrinsically fluorescent, no chemical tagging or substrate addition is necessary to generate the optical signal. Here this approach was used to develop a biosensor of receptor-arrestin interaction (Fig. 1a). β-arrestin2 and the fluorescent protein mNeonGreen ^37,38^ were fused together such that the entire arrestin molecule was inserted into the critical seventh stave of mNeonGreen (see Methods).

**Figure 1.**
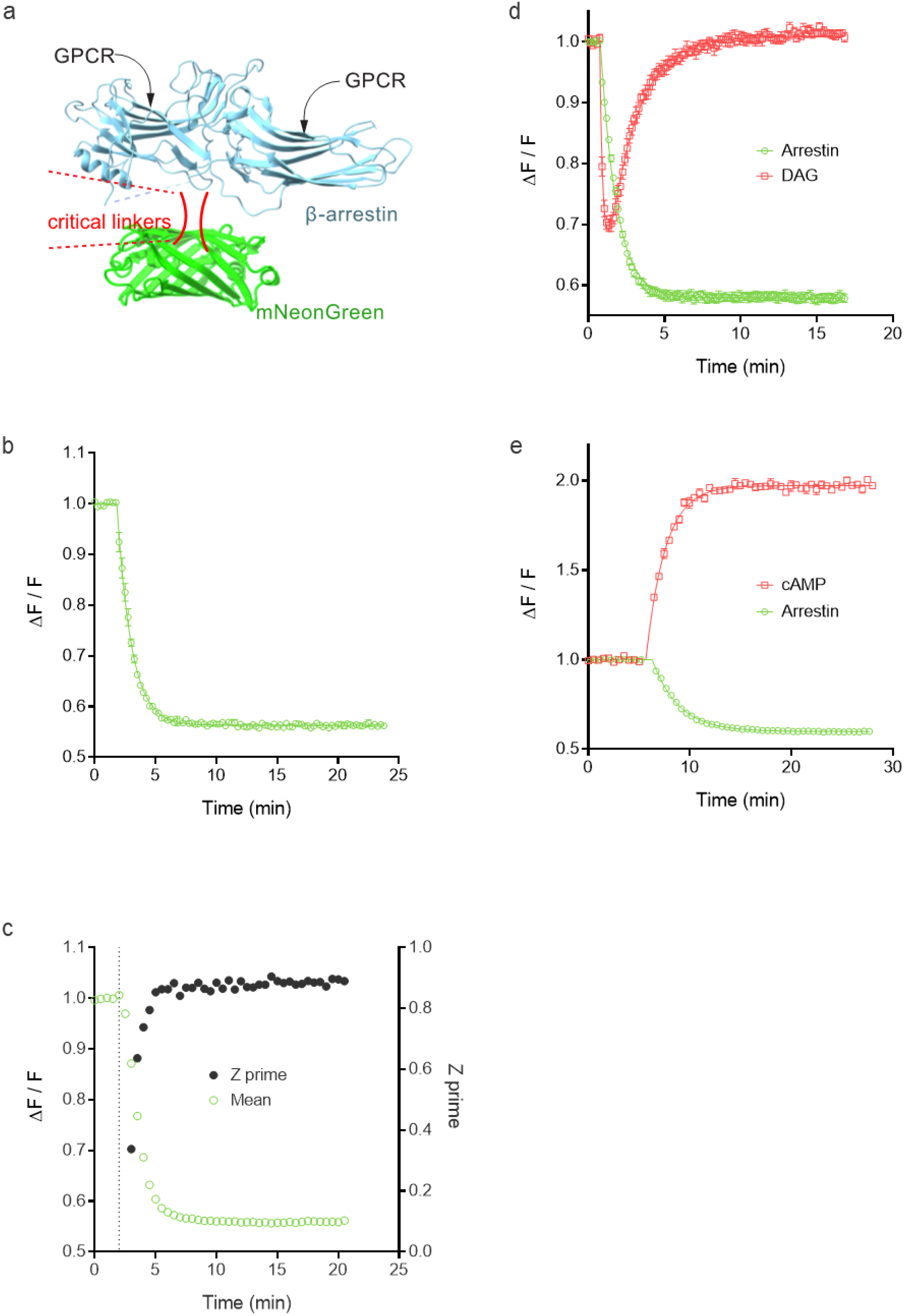
Characterization of a new arrestin sensor, (a) Schematic illustration of fluorescent arrestin biosensor. β-arrestin2 was engineered to incorporate mNeonGreen using a screening process to identify optimal positioning and linking for generation of fluorescent signal, (b) Change in arrestin fluorescence on activation of the angiotensin AT_1_ receptor by Angll at a concentration of 32 μM. Receptor-arrestin interaction results in a decrease of sensor fluorescence intensity, (c) Z’ values over the time course of AT_1_ receptor-arrestin interaction stimulated by lOpM Angll. (d) Multiplexing green arrestin sensor with red diacylglycerol sensor, with activation of the AT_1_ receptor by 30 μM Angll. (e) Multiplexing with red cADDIs (cAMP) sensor, with activation of the V_2_ vasopressin receptor by vasopressin at 30 μM. Data was generated with the BMG CARIOstar (b-d) or Biotek Synergy Mx (e) plate readers. Data points are mean ± sem [n = 2 for (b) and n = 4 for (d) and (e)].

Agonist application resulted in a robust change of fluorescence intensity of the arrestin sensor, for the angiotensin AT_1_ receptor (Fig. 1b) and V_2_ vasopressin receptor (Fig. 1e). The signal was quantified as the fluorescence after agonist addition normalized to that of the baseline signal before addition (ΔF / F). The robustness of the signal over time is a key determinant of utility for kinetic application of the sensor. Consequently, statistical analysis was conducted for the ΔF / F value at all time points measured. The angiotensin data was used for this purpose. The coefficient of variance (% CV) of the technical replicates (duplicates) is shown in Supplementary Fig. S1, for multiple concentrations of AngII spanning the effective concentration range. The % CV was less than 7% for all data points, and less than 5% in 98% of cases. This result indicates the signal is sufficiently robust to quantify the signal over the entire time course across the effective concentration range. Not surprisingly, the error was greatest when the signal was changing the most over time, i.e. on the linear part of the curve (Supplementary Fig. S1). Next we assessed the Z’ value over time to determine the ideal timeframe for a single time point measurement (Fig. 1c), the paradigm typically used for high throughput screening (HTS). Z’ is lowest on the rise phase of the time course, when the signal is changing the most over time and the magnitude of the signal is low relative to the plateau phase (Fig. 1c). At the plateau, the Z’ value was high and consistent over time. These findings indicate a time point at the plateau phase is ideal for HTS and that the signal for the AT_1_ receptor is potentially robust enough for HTS.

We tested whether the arrestin sensor could be multiplexed with sensors of G-protein signaling, i.e. that the signals could be detected in the same well. The green arrestin sensor could be combined with the red diacylglycerol sensor (Fig. 1d, AT_1_ receptor) and red cAMP sensor (Fig. 1e, V_2_ receptor). This capability allows direct comparison of the kinetics of arrestin recruitment and G-protein signaling. For both the AT_1_ receptor and V_2_ receptor, the recruitment of arrestin occurs within the timeframe of the attenuation of the G-protein-mediated signal (decline of the DAG signal (Fig 1d) and approach to plateau of cAMP concentration (Fig. 1e)). This finding is consistent with arrestin recruitment regulating (attenuating) G-protein signaling via these receptors.

### Time course and concentration-dependence of arrestin recruitment by the angiotensin AT_1_ receptor

We next characterized the arrestin recruitment kinetics of the AT_1_ receptor, examining the shape of the time course and the concentration-response characteristics. For the full agonist and endogenous ligand AngII, arrestin was initially recruited rapidly at a maximally-effective concentration (32 μM), starting within 1 minute of application (Fig. 2a). The recruitment leveled off then reached a plateau within five minutes (Fig. 2a). The plateau was stable for the remainder of the measurement period (for example, 20 min, Fig. 2a), indicating a steady-state had been obtained. By visual inspection, this profile appeared to conform to the familiar association exponential curve. The data were fit to this equation using Prism 8.0:

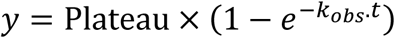

where Plateau is the response at the asymptote (at infinite time) and *k*_obs_ the rate constant. Data were fit well by this equation (R^2^ correlation coefficient > 0.99 in all cases). The *t*_1/2_, calculated from *k*_obs_ (*t*_1/2_ = 0.693/ *k*_obs_) was 44 ± 3 sec at 32 μM (Supplementary Table 1). For lower, sub maximally-effective concentrations, e.g. 32 nM, the initial recruitment was slower, manifest as a shallower initial rise, and the plateau was lower (Fig. 2a). Data for lower concentrations were also fit well by the association exponential equation (Fig. 2a, R^2^ > 0.95 in all cases). The concentration-dependence of the Plateau and *k*_obs_ parameter values is shown in Supplementary Fig. S2 - both Plateau and *k*_obs_ increased as the AngII concentration was increased.

**Figure 2.**
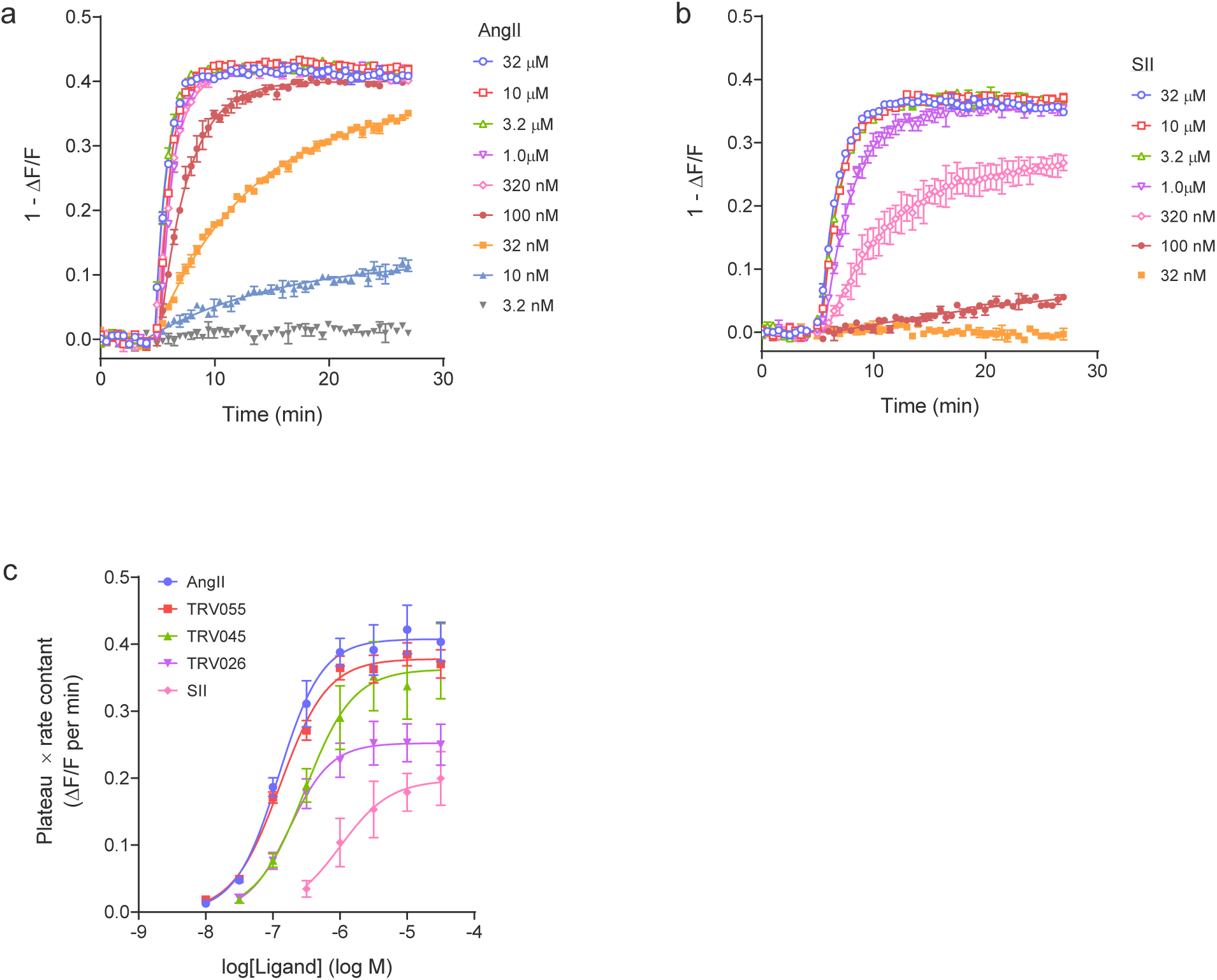
Dose response kinetic analysis for arrestin recruitment to the AT_1_ angiotensin receptor. The time course of the arrestin sensor response was measured for a range of concentrations of AT_1_ receptor agonists AngII (a), SII (b) and TRV055, TRV045 and TRV026 (Supplementary Fig. S6). Data are well fit by the association exponential equation, 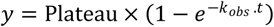, as predicted by the kinetic model of arrestin recruitment (Appendix). From these fits it is possible to measure *k*_*τ*_, the initial rate of arrestin recruitment by the agonist-bound receptor. First, the Plateau value is multiplied by the *k*_*obs*_ value. The resulting value is plotted against the agonist concentration, shown in panel (c). These data are then fit to a dose-response equation (“Log(agonist) vs. response -- Variable slope” equation in Prism ^16^). *k*_*τ*_ is the top of the curve, the Plateau ×*k*_obs_ value at maximally-effective agonist concentrations. Agonist affinity for the receptor is the EC_50_ of the sigmoid curve. Data are from the Biotek Synergy Mx plate reader.

A known partial agonist for arrestin recruitment was then tested, [sarcosine^1^,Ile^4^,Ile^8^]AngII (SII) ^39^. Superficially, visual inspection of the plots suggests little difference between SII and AngII at the maximally-effective concentration of 32 μM (Fig. 2a and b). However, the ability to accurately quantify the kinetics of the response indicated an appreciable difference in the rate of arrestin recruitment; the *t*_1/2_ of SII calculated from *k*_obs_ was 84, approximately twice that of AngII (45 sec) (Supplementary Table S1). This indicated SII recruits arrestin more slowly than AngII, providing a kinetic perspective on its partial agonist activity. By contrast, at the plateau, there was no appreciable difference between SII and AngII (compare 32 μM data in Fig. 2a and b, Supplementary Table S1). This finding indicates that if arrestin recruitment was measured at a single time point on the plateau, the difference of activity between the peptides would not have been detected; SII would have appeared to be a full agonist. Only by measuring the rate of arrestin recruitment was the partial agonist activity revealed.

### Pharmacological analysis model of receptor-arrestin interaction kinetics

Applying kinetics to development of new ligands in pharmacological discovery requires the extraction of simple drug parameters from time course data. The time course of arrestin recruitment was conformed to an association exponential curve for the AT_1_ receptor (see above) and the V_2_ receptor (Fig. 1e). A number of empirical drug parameters can be obtained from these data, such as the *t*_1/2_, plateau, AUC, or the signal at a single time point. An alternative and biologically meaningful parameter is the initial rate of signaling, analogous to the initial rate of enzyme activity ^24,25^. We recently discovered this parameter can be easily obtainable from signaling kinetic data using a new kinetic pharmacological framework of GPCR signaling (Fig 3) ^25^. This approach is based on the principles of enzyme kinetic data analysis. An enzyme converts a substrate into a product (Fig. 3a). By analogy, a GPCR converts a precursor of the signal into the signal (Fig. 3b), for example, conversion of inactive G-protein to active G-protein. For enzymes, the initial rate of activity is the rate before it becomes limited by regulation mechanisms and depletion of the substrate. By analogy, the initial rate of GPCR signaling is the rate before signaling regulation mechanisms limit the signal (Fig 3c) ^25^. The canonical short-term regulation of signaling mechanisms are receptor desensitization, and degradation of the signal (e.g. hydrolysis of GTP bound to G-protein or clearance of cytoplasmic Ca^2+^). The response can also become limited by depletion of the precursor of the signal, e.g. depletion of Ca^2+^ from intracellular stores. The initial rate is defined by the law of mass action, being a function of the interacting components and a microscopic rate constant. For enzymes the initial rate is [*E*]_*TOT*_[*S*]_*TOT*_*k*_*CAT*_(product of total enzyme, total substrate, and the catalytic rate constant). By analogy, for GPCR signaling, the initial rate is *E*_*P(TOT)*_[*R*]_*TOT*_*k*_*E*_, where *EP(TOT)* is the total precursor, [*R*]_*TOT*_ the total receptor concentration, and *k*_*E*_ a rate constant termed the transduction rate constant. This is the initial rate of signaling by the ligand-bound receptor and is termed *k*_*τ*_. This parameter can be easily estimated by curve fitting ^24,25^.

**Figure 3.**
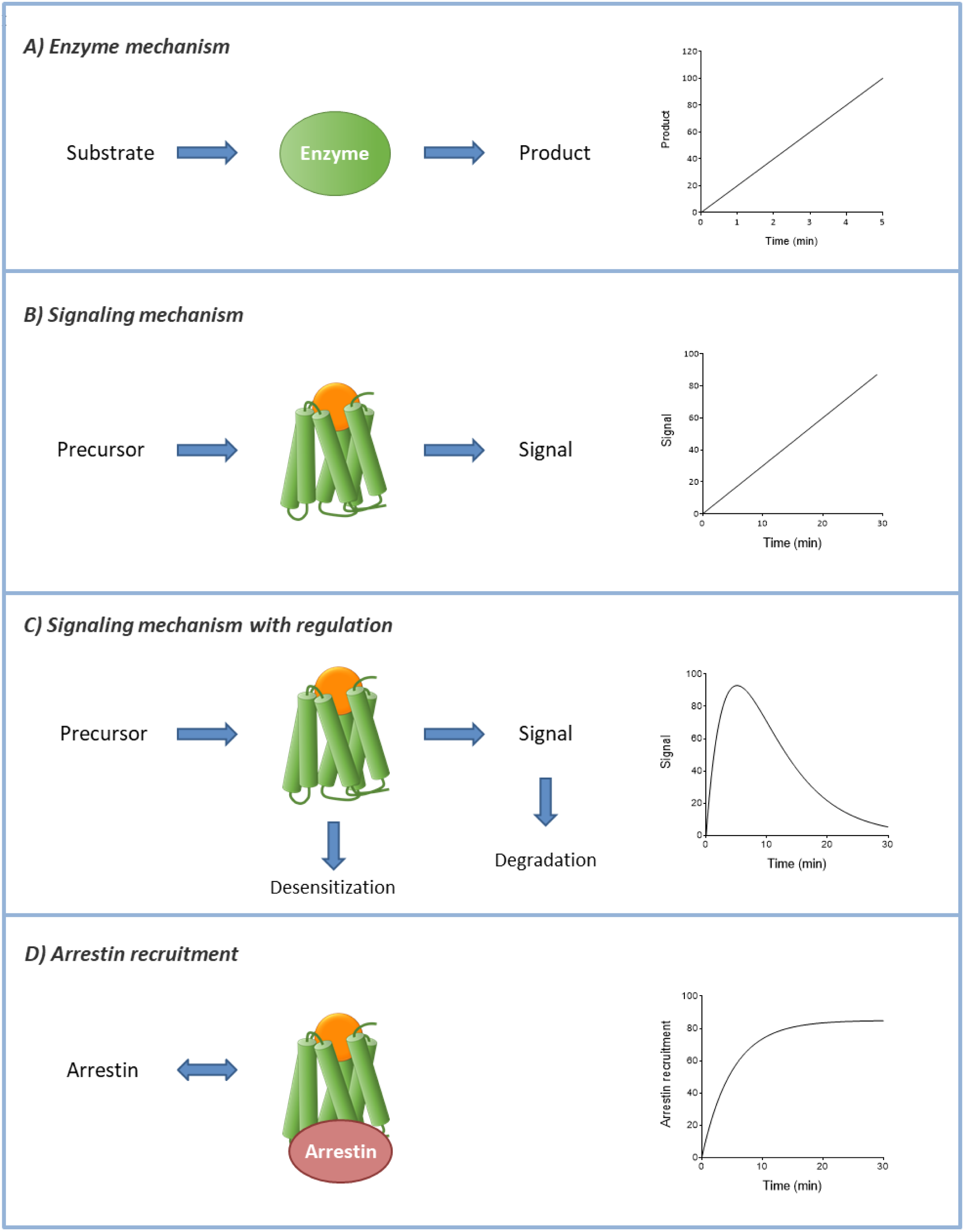
Receptor signaling kinetics mechanisms. A data analysis framework for quantifying the kinetics of GPCR signaling has been developed previously ^24,25^. This method quantifies the initial rate of signaling (*k*_*τ*_), analogous to the initial rate of enzyme activity. Here a new analysis method is introduced to quantify arrestin recruitment kinetics, (a) Enzyme catalysis mechanism - enzyme converts substrate to product. The time course is a straight line (under conditions of minimal substrate depletion), (b) GPCR signaling mechanism - agonist-bound GPCR converts a signal precursor to the signal (e.g. GDP-bound G-protein to the GTP-boiuid form, or sequestered Ca^2+^ to cytoplasmic Ca^2+^). The time course is a straight line, (c) GPCR signaling modulated by regulation of signaling mechanisms - receptor desensitization and signal degradation. The resulting time course is a rise-and-fall curve. The shape of the time course is dependent the number and nature of regulation mechanisms^25^. (A third regulation mechanism is depletion of signal precursor.) (d) Arresgin recruitment - agonist-bound GPCR interacts with arrestin. The time course is an association exponential curve, as described in Results and Appendix.

Here we developed a pharmacological analysis that can be applied to measure the initial rate of arrestin recruitment, the direct analogue of the initial rate of signaling described above. The mechanism of arrestin recruitment is known. Agonist-bound GPCR is phosphorylated by kinase enzymes ^40,41^ and the phosphorylated receptor binds arrestin ^42,43^. This mechanism is represented as follows:

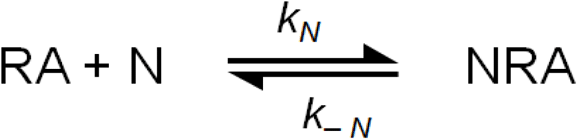

Arrestin (N) interacts with ligand-bound receptor (RA), governed by the rate constant *k*_*N*_. (In most cases the rate-limiting step in arrestin recruitment is receptor phosphorylation so in these cases *k*_*N*_ is the rate constant for receptor phosphorylation). Arrestin dissociates from the ligand-receptor complex, governed by the arrestin-receptor dissociation rate constant *k*_*-N*_. In the Appendix an equation was derived that describes the level of ligand-receptor-arrestin complex over time after the addition of ligand (equation (3)).

The validity of the model was assessed by comparing data simulated using the model with experimental data. In agreement with experimental data, the simulated arrestin recruitment conforms to an association exponential curve (Supplementary Fig. S3a). This is because equation (3) is of the form of the association exponential equation (Appendix). The concentration-response was simulated (Supplementary Fig. S3a) and the resulting *k*_obs_ and Plateau value of the association exponential fit determined (Supplementary Fig. S3b,c). Increasing the agonist concentration increased Plateau and *k*_obs_, with both effects approaching a limit at maximally-effective agonist concentrations (Supplementary Fig. S3). This profile was in agreement with the experimental concentration-response data for the AT_1_ receptor (Supplementary Fig. S2).

*k*_*τ*_ for arrestin recruitment by the ligand-bound receptor is a measurable parameter in the model. This parameter is [N]_TOT_[R]_TOT_*k*_*N*_, where [N]_TOT_ is total arrestin concentration. It emerges that this parameter can be quantified using the Plateau and *k*_obs_ parameters from the fit to the association exponential equation (Appendix). Specifically, *k*_*τ*_ is equal to the Plateau value multiplied by the *k*_obs_ value at a maximally-stimulating concentration of ligand. This can be determined using either a full concentration response (Supplementary Fig. S4) or just a maximally-stimulating concentration (Supplementary Fig. S5), as described below.

### Quantifying arrestin recruitment kinetics for the angiotensin AT_1_ receptor - concentration-response

The angiotensin AT_1_ receptor is a prototypical GPCR in the study of arrestin recruitment and biased signaling. The receptor for angiotensin II (AngII), the AT_1_ receptor, regulates blood pressure and consequently is a target for antihypertensive drugs (the sartan antagonists) ^44^. Biased ligands at the AT_1_ receptor that selectively promote arrestin signaling while blocking G-protein signaling can elicit increased cardiac performance compared with unbiased ligands, potentially beneficial for treating cardiovascular disorders ^28-31^.

We used the *k*_*τ*_ method to quantify the kinetics of arrestin recruitment of five AT_1_ receptor ligands with known varying degrees of arrestin recruitment and bias. The method for quantifying *k*_*τ*_ is as follows (illustrated schematically in Supplementary Fig. S4). First, the time course data for the effective concentrations (10 nM - 32 μM for AngII) were fit to the association exponential equation (Fig. 2a). From these fits the fitted parameter values were taken for Plateau and *k*_obs_. These values were then multiplied together. The resulting Plateau x *k*_obs_ values were then plotted against the ligand concentration, as shown in Fig. 2c. The resulting plot was a sigmoid curve (consistent with the theoretical prediction of the model (Supplementary Fig. 3d)). The data were then fit to the sigmoid curve equation, for example the “log(agonist) vs. response -- Variable slope” equation in Prism ^45^. From this fit *k*_*τ*_ was obtained - it is the value of the asymptote. More precisely, *k*_*τ*_ is the Plateau x *k*_obs_ value for a maximally-effective concentration of ligand. The theoretical basis for this calculation is shown in Appendix, “Defining the initial rate of arrestin recruitment and identifying in the equations.” The fitted value for *k*_*τ*_ for AngII was 0.41 ± 0.03 normalized fluorescence units (NFU) per min (Table 1).

**Table 1:** Arrestin dose response parameters from the kinetic model applied to the AT_1_ angiotensin receptor

| Ligand | $k_{\tau}$<br>(NFU.min<sup>-1</sup>)<sup>1</sup> | $k_{\tau}$<br>(% AngII) | $\text{p}K_A$ | $K_A$<br>(nM) |
| --- | --- | --- | --- | --- |
| AngII | 0.41 ± 0.03 | 100 | 6.92 ± 0.06 | 120 |
| TRV055 | 0.38 ± 0.02 | 93 | 6.90 ± 0.04 | 130 |
| TRV045 | 0.36 ± 0.05 | 89 | 6.52 ± 0.02 | 300 |
| TRV026 | 0.25 ± 0.03 | 62 | 6.76 ± 0.05 | 180 |
| SII | 0.20 ± 0.03 | 48 | 5.97 ± 0.15 | 1,100 |

We next evaluated four synthetic AT_1_ receptor ligands, SII, TRV120055, TRV120045 and TRV120026 ^39,46^. (For clarity, the name of the last three compounds is abbreviated to TRV055, TRV045 and TRV026). These compounds were developed in SAR campaigns aiming to identify biased ligands for the AT_1_ receptor. Applying the kinetic analysis method to the SII data gave the Plateau x *k*_obs_ vs ligand concentration data in Fig 2c. Fitting the sigmoid curve equation to these data gave a *k*_*τ*_ value of 0.20 NFU per min (Table 1). This value is approximately half that of AngII (0.41 NFU per min). This means the initial rate of arrestin recruitment by the SII-receptor complex is approximately half that of the AngII-receptor complex. Note this provides a biologically meaningful kinetic scaling of the partial agonist activity of SII for arrestin recruitment. The degree of partial agonism can be quantified conventionally, by dividing *k*_*τ*_ of SII by that of AngII. This gave a normalized *k*_*τ*_ value of 48 % for SII (Table 1).

This method was applied to the remaining three ligands. In all cases the time course data conformed to the kinetic model, being well-fitted by the association exponential equation (Supplementary Fig. S6) with the Plateau and *k*_obs_ values being dependent on agonist concentration (Supplementary Fig. S2). Applying the kinetic analysis method gave the sigmoid curves in Fig 2c and the *k*_*τ*_ values in Table 1. TRV055 and TRV045 are effectively full agonists for recruiting arrestin at the initial rate (*k*_*τ*_ 93% and 89%, respectively, of that of AngII). The efficacy of TRV026 (62%) was intermediate between that of TRV055 and SII (Fig. 2c, Table 1).

The analysis also provides an estimate of ligand affinity for the receptor, *K*_*A*_. This is given by the EC_50_of the Plateau x *k*_obs_vs concentration sigmoid curve (Appendix, Supplementary Fig. S4). With the exception of SII, the affinity of the ligands was similar (120 - 300 nM, Table 1). The affinity of SII was lower (1,100 nM, Table 1).

### Quantifying arrestin recruitment kinetics for the angiotensin AT_1_ receptor - single concentration

A simplified method is feasible for measuring *k*_*τ*_ (see Supplementary Fig. S5). All that is required is a time course of response measurement at a single, maximally-stimulating concentration of ligand (maximally-stimulating at all time points). In Fig. 4a the data for the maximally-effective concentration of the AT_1_ receptor peptides is presented (32 μM). The data were fit to the association exponential equation to determine Plateau and *k*_obs_. *k*_*τ*_ was then calculated - it is the Plateau multiplied by *k*_obs_ for a maximally-stimulating concentration of ligand, as explained in the Appendix. The *k*_*τ*_ values for the five ligands tested are shown in Table 2 and are in good agreement with the values determined using the concentration-response method (Table 1).

**Table 2.** *k*_*τ*_ values and ratios for AT_1_ angiotensin receptor-mediated arrestin recruitment and calcium mobilization

| Ligand | Arrestin | | Diacylglycerol | | Arrestin/DAG<br>$k_\tau$ ratio<br>(% AngII ratio) | Calcium | | Arrestin/$\text{Ca}^{2+}$<br>$k_\tau$ ratio<br>(% AngII ratio) |
| --- | --- | --- | --- | --- | --- | --- | --- | --- |
| | $k_\tau$<br>(NFU.min<sup>-1</sup>)<sup>1</sup> | $k_\tau$<br>(% AngII) | $k_\tau$<br>(NFU.min<sup>-1</sup>)<sup>1</sup> | $k_\tau$<br>(% AngII) | | $k_\tau$<br>(NFU.min<sup>-1</sup>)<sup>1</sup> | $k_\tau$<br>(% AngII) | |
| AngII | 0.40 ± 0.03 | 100 | 1.7 ± 0.2 | 100 | 1.00 | 2.2 ± 0.2 | 100 | 1.0 |
| TRV120055 | 0.37 ± 0.02 | 92 | 2.1 ± 0.1 | 120 | 0.74 | 2.5 ± 0.2 | 110 | 0.81 |
| TRV120045 | 0.38 ± 0.06 | 93 |  |  |  | 0.66 ± 0.02 | 30 | 3.1 |
| TRV120026 | 0.25 ± 0.03 | 62 |  |  |  | 0.46 ± 0.08 | 21 | 3.0 |
| SII | 0.20 ± 0.04 | 49 |  |  |  | 0.25 ± 0.06 | 12 | 4.2 |

**Figure 4.**
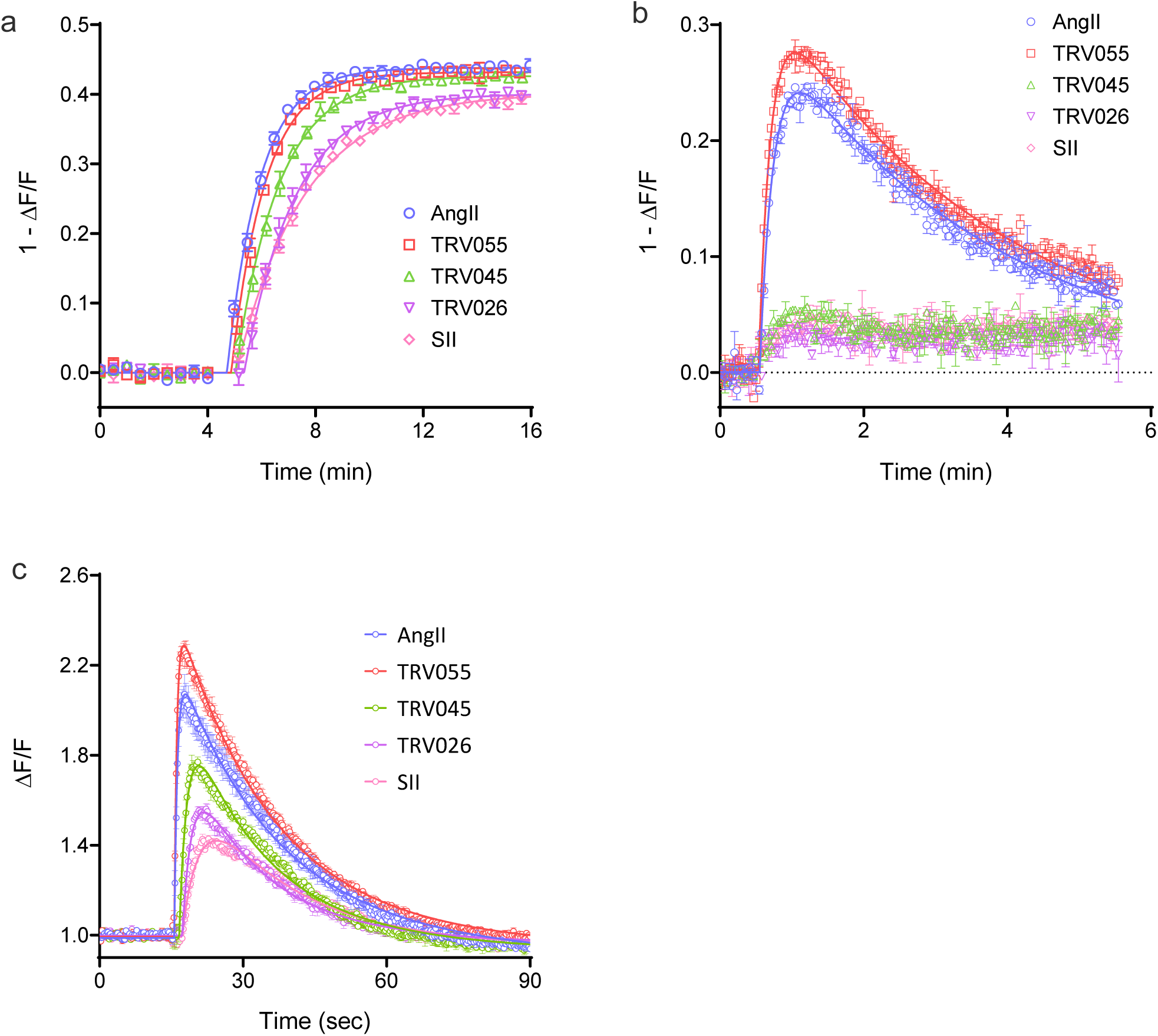
Kinetics of arrestin recruitment and G-protein signaling via the AT_1_ angiotensin receptor, analyzed using the kinetic model. Five ligands with known signaling bias were tested for arrestin recruitment (a), DAG production (b) and Ca^2+^ mobilization (c). A maximally-stimulating concentration of ligand was used (32 µM), enabling *k*_*τ*_ to be quantified as described in Supplementary Fig. S5 for arrestin recruitment. For DAG production and Ca2+ mobilization *k*_*τ*_ was determined by fitting to the rise-and-fall exponential equation 24,25 (see Methods); the fitted value of *C* is equal to *k*_*τ*_. The *k*_*τ*_values are given in Table 2. Note the signal for arrestin and DAG has been normalized to give an upward response to the downward sensor (1 - ΔF/F). All data are from the Biotek Synergy Mx plate reader.

### Application to quantifying biased agonism

Biased agonism quantification relates the capacity of a ligand to activate one pathway relative to one or more other pathways ^9-12^. Numerous methods and scales have been developed and successfully applied ^8,46-53^. One approach is to compare ligand efficacy, i.e. the capacity of the agonist-occupied receptor to generate the signal. *k*_*τ*_ provides such an efficacy value, being the initial rate of signal generation by the agonist-occupied receptor. Here *k*_*τ*_ is used to assess signaling bias of the AT_1_ receptor ligands described above. These ligands have been reported to vary in their bias, with SII, TRV026 and TRV045 being arrestin biased ^54,55^ and TRV055 being unbiased ^46^ relative to AngII ^46^. In this study, G-protein and arrestin signaling was measured using the same fluorescent biosensor assay modality, enabling near-identical conditions to be employed in comparing the pathways. The Gq pathway was quantified at the level of diacylglycerol using the Red Downward DAG sensor ^21^ (Fig. 4b), and, one step downstream, Ca^2+^ mobilization measured using the R-GECO sensor ^22^ (Fig. 4c).

The β-arrestin, DAG and Ca^2+^data are shown in Fig. 4, for a maximally-stimulating concentration of the five ligands (32 μM). It is immediately obvious that curve shapes are different for the pathways - an association exponential curve defines arrestin recruitment (Fig. 4a) and a rise-and-fall curve defines DAG and Ca^2+^ signaling (Fig. 4b and c). The kinetic analysis method was designed to handle this scenario - the initial rate can be extracted from different curve shapes ^24,25^. The initial rate of arrestin recruitment was quantified as described above, by multiplying the rate constant by the plateau of the association exponential fit, and the resulting *k*_*τ*_ values are in Table 2). For DAG and Ca^2+^, a rise-and-fall exponential equation was applied:

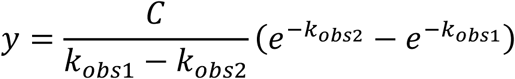

*k*_*τ*_ is equal to the value of *C* in this equation ^24,25^ The equation assumes two regulation mechanisms are in operation, corresponding to the two exponent terms. (These are most likely receptor desensitization and response degradation for DAG, and precursor depletion and response degradation for the shorter-term Ca^2+^ response ^24^). The fitted *k*_*τ*_ values for DAG and Ca^2+^ are shown in Table 2. For DAG, no response was detectable for TRV045, TRV026 and SII. Responses to these peptides were observed for Ca^2+^, one step downstream and so potentially more sensitive to small effects owing to signal amplification.

The next step in the bias calculation is normalization to a reference agonist. AngII was used for this purpose, being a full agonist and the endogenous ligand. The resulting normalized *k*_*τ*_ values are shown in Table 2. A qualitative assessment of bias can be made with these values. All ligands appreciably stimulate arrestin recruitment (by ≥ 49%, Table 2). By contrast, only weak activation of the Gq pathway was detected for three of the ligands (TRV045, TRV026 and SII); DAG was not detectable, and only a weak maximal effect was observed in the more amplified Ca^2+^ signal (12-30 %, Table 2). This finding suggests the ligands are biased for arrestin over Gq signaling.

The final, quantitative step in the bias calculation is the bias ratio calculation. This was done by dividing the normalized *kτ* value for arrestin recruitment by that for Ca^2+^. By definition, the ratio for the reference ligand AngII was 1.0 (Table 2). Three ligands were arrestin-biased, evident by their arrestin/Ca^2+^ *k*_*τ*_ ratio being greater than 1 (3.1, 3.0 and 4.2 for TRV045, TRV026 and SII, Table 2). One ligand was unbiased, relative to AngII (TRV055, Table 2). These results are consistent with the known bias profiles of these ligands ^46,54,55^. For example, SII is an established arrestin-biased AT_1_ receptor ligand, and TRV055 is known as a balanced ligand (relative to AngII). In Fig. 5, the bias factor from the *k*_*τ*_ ratio was compared with the bias factor calculated previously using a validated method, the operational model approach ^46^. The bias profile across the four ligands tested was similar; TRV12055 was similar to AngII, whereas TRV045 and TRV026 were arrestin-biased and the degree of bias was similar for both ligands (Fig. 5).

**Figure 5.**
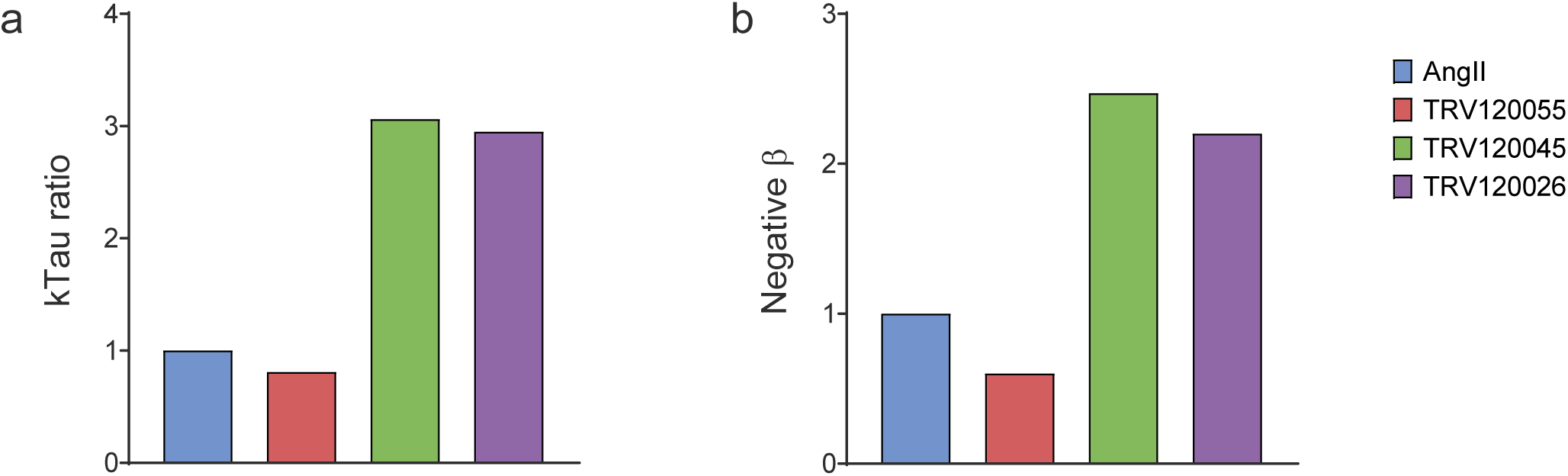
Comparison of *k*_*τ*_ bias ratio with published bias values obtained using the operational model. Bias for arrestin recruitment over Gq signaling via the AT1 receptor are shown. The *k*_*τ*_ bias ratio for B-arrestin-2 recruitment over Ca^2+^ mobilization (A) was calculated as shown in Table 1. In (b) the bias factor was obtained using the operational model applied to data at a single time point, for beta-arrestin-2 recruitment and inositol-1-phosphate production. Data are from Table 2 of ref 44.

These results indicate bias can be calculated simply and in kinetic terms using a common data analysis framework, the *k*_*τ*_ method, for G-protein signaling and arrestin recruitment. The application of a common detection modality for arrestin and G-protein (the genetically-encoded sensors) also provides a unified technical platform that simplifies the interpretation of bias.

## DISCUSSION

Arrestin recruitment to GPCRs is the first step in a signaling pathway that mediates myriad GPCR responses ^26,27^ that are often selectively activated over G-protein pathways by biased agonists ^10,15,36,54,55^. Here we developed a new method of quantifying arrestin interaction that takes into account the kinetics, i.e. the time course, of the response. We developed an improved kinetic arrestin assay. Present assays include endpoint assays, requiring a plate for every time point, or employ bioluminscent resonance energy transfer-based sensors that often require read times too long to accommodate the rapid changes of arrestin recruitment typically observed for GPCRs. A new data analysis framework was developed to analyze time course data to quantify arrestin recruitment in kinetic terms, specifically the initial rate of recruitment. The resulting values can be directly compared with the initial rate of signaling through other pathways, enabling straightforward assessment of biased agonism. Application of this approach to biased agonists of the AT_1_ angiotensin receptor provided bias estimates similar to literature values, suggesting the method can be applied generally to quantify biased agonism.

The β-arrestin sensor is a genetically-encoded arrestin protein modified to incorporate the intrinsically-fluorescent protein mNeonGreen ^37,38^. The fluorescent protein is incorporated in such a way as to render the fluorescence emission conformationally-sensitive, such that interaction with the GPCR results in a decrease of fluorescence intensity. The sensor was designed to meet criteria necessary for application to drug discovery. The high quantum yield provides a large enough signal for detection in plate readers and enables short excitation times to be used, which minimizes photobleaching ^21^. BacMam provides consistent receptor and biosensor expression, minimizing cell to cell and well to well variability. Importantly the BacMam delivery can adjust to optimize expression levels. The Z-score was high enough for the AT_1_ and V_2_ receptors for application to HTS and lead optimization. Minimal steps are required for the assay; once the cells are prepared, the only reagent addition step is the application of agonist. The absence of subsequent detection reagent additions improves workflow and likely contributes to the high Z-score. The disadvantage of the sensor for HTS is that fluorescent compounds can potentially interfere with the assay because direct fluorescence excitation-emission is employed, requiring a follow-up assay to test for compound fluorescence. This is likely less of an issue in lead optimization, by which time fluorescent molecules, usually undesirable, have been disregarded. The sensor also possesses desirable pharmacological properties that simplify the analysis and interpretation of data in lead optimization, specifically, 1) The unmodified receptor can be used, providing estimates of authentic rather than forced coupling to arrestin. 2) The detection of the interaction is at the level of the receptor interaction itself, rather than downstream (for example, activation of transcription factors or phosphorylation of extracellular signal-regulated kinase). As a result, there is no receptor reserve, simplifying interpretation of ligand efficacy for arrestin recruitment; the maximal response to ligand is equal to the efficacy of the ligand. The sensor is ideal for kinetic measurements. The short excitation time enables a short time interval between data points of the time course (e.g. 9 sec, Fig. 1d). The high reproducibility provides robust data at early time points, when the change of signal is small but the value of the data point to defining the kinetics is high (Fig. 1c, Supplementary Fig. 1). Finally, two or more pathways can be quantified using the same modality (and in the case of multiplexing, in the same cells, Fig. 1d,e), ideal for assessing biased agonism. This avoids complications arising from the use of different assay systems for the pathways being compared, for example different biological material (e.g. membranes for GTPγS binding and cells for arrestin recruitment), different time points, buffer conditions, temperatures and so on.

A data analysis framework was required to translate the arrestin time course data into a useful pharmacological parameter. Previously we have quantified G-protein signaling kinetics using the initial rate of signaling, which we termed *k*_*τ*_ ^24,25^. Here this concept was extended to arrestin recruitment. (This was necessary because the original mechanism described generation of a downstream signal, whereas the arrestin sensor signal reports direct receptor-effector interaction.) The resulting analysis quantifies the initial rate of arrestin recruitment, i.e. the same parameter as that used previously for G-protein signaling. *k*_*τ*_ is reasonably straightforward to measure from time course data, using either a full concentration-response (Fig. 2c, Supplementary Fig. S4) or at a single maximally-effective concentration (Fig. 4a, Supplementary Fig. S5). Time course data are fitted to the association exponential equation, fitted values of Plateau and *k*_obs_ obtained, then these values multiplied together. *k*_*τ*_ is Plateau x *k*_obs_ at a maximally-effective ligand concentration.

The *k*_*τ*_ parameter has certain benefits as a pharmacological parameter. First, it takes into account the kinetics of signaling, being a rate. In simple terms, *k*_*τ*_ is the same at all time points. This avoids the problem of time-dependence of ligand efficacy values that can occur when using a single time point assay ^4,8^. Second, the initial rate is biologically meaningful; it describes the response generation (arrestin recruitment or G-protein signaling) by the receptor before the response becomes modulated by regulation of signaling mechanisms (e.g. dissociation of arrestin, and degradation of second messengers). For this reason *k*_*τ*_ is potentially more intuitive to use in interpreting and translating ligand efficacy than more abstract pharmacological parameters such as τ in the operational model. However, the *k*_*τ*_ method is relatively new and so not established. Currently unexplored is the impact of receptor reserve on *k*_*τ*_ estimates of downstream signaling. The current theoretical framework assumes *k*_*τ*_ incorporates receptor reserve ^24^ but the extent to which this is sufficient to explain experimental data remains to be determined.

This analysis was applied to biased agonism assessment using the AT_1_ receptor (Fig. 4, Table 2). Bias between arrestin and Gq-mediated signaling was quantified using the *k*_*τ*_ method for five ligands. Bias was quantified as the ratio between the *k*_*τ*_ value, normalized to that of AngII, for arrestin and Ca^2+^. The resulting bias estimates were in good agreement with those from previous studies (Fig. 5) ^46^, validating the method. The new sensor and analysis platform provides an ideal system for measuring biased agonism. Technical differences between signaling assay readouts are minimized because the same assay conditions are employed for arrestin and G-protein signaling measurements - the only difference is the sensor introduced into the cells the day before assay. [Though not used in this study, the sensors can be multiplexed, which eliminates assay condition differences completely (Fig. 1d,e).] The use of the kinetic paradigm and *k*_*τ*_ eliminates the time dependence of biased agonism estimates that can result from the use of endpoint assays and single time point data analysis ^8^. The unified data analysis framework enables arrestin and G-protein signaling to be directly compared using the same parameter (initial rate of arrestin recruitment or signal generation). The new analysis somewhat simplifies the determination of the bias factor. The current analysis methods are quite complex ^8,46-53^, involving multiple calculations and in some cases advanced curve fitting (such as simultaneous fitting of multiple ligand concentration-response curves). The *k*_*τ*_ method is somewhat simpler, requiring a single bias calculation (the *k*_*τ*_ ratio) and employing basic, familiar curve fitting. Finally, the bias scale is more straightforward; the bias scales of existing methods are abstract, whereas the *k*_*τ*_ ratio is a more biologically meaningful parameter, being the ratio of the initial rate of the responses being compared.

In summary, in this study we describe a new platform that utilizes the kinetics of response to quantify arrestin recruitment and G-protein-mediated signaling. This provides a universal analysis framework employing the initial rate of activity that simplifies quantification and interpretation of ligand activity and biased agonism. This method can be employed by drug discovery scientists to improve the identification, optimization and development of new therapeutics.

## METHODS

### Sensor Design

The design goals for a new β-arrestin2 sensor were: 1) it needed to be bright enough for detection on automated fluorescence plate readers, 2) it needed to use a single fluorescent protein so that other portions of the visible spectrum were available for other colored sensors and multiplex recordings, and 3) the change in fluorescence in response to GPCR activation has to be large enough to produce signal-to-noise ratios of > 60.

Using the β-arrestin2 structures as our guide, we designed a series of constructs in which the entire β-arrestin2 was inserted into the middle of the seventh stave of beta sheet in the barrel of mNeonGreen, directly adjacent to the chromophore ^37,38^. The goal was to convert the change in β-arrestin shape when it binds a phosphorylated receptor into a change in the chromophore environment resulting in a change in fluorescence intensity. The first biosensors that positioned analyte binding domains in this position used a circularly permuted version of the fluorescent protein, and include the GCaMP and GECO Ca^2+^ sensors, the upward and downward DAG sensors, and the caDDis cAMP sensor ^21^. More recently it has become clearer that if the termini of the analyte binding domain are close to one another, then the entire binding domain can be inserted into the 7th stave of the fluorescent protein without having to circularly permute the fluorescent protein ^56^.

The initial constructs were screened for responses on a fluorescence microscope with time lapse imaging, and drugs were added to the well by hand. HEK 293T cells were transiently transfected with the sensor prototypes as well the AT_1_ angiotensin receptor. The receptor was then activated with the addition of 30 μM AngII, and the sensor was monitored for changes in fluorescence. After identifying a functional prototype, mutagenic PCR was used to randomly mutagenize two to three amino acids at a time at the fusion junction/s, producing random libraries of thousands of mutants. These mutants were then screened in a high throughout format on a fluorescence plate reader for AT_1_ receptor activation-dependent changes in fluorescence intensity.

### BacMam production and titer

BacMam was produced following the methods described previously ^21^. To establish BacMam titers, we quantified viral genes per mL using quantitative PCR. Samples are diluted 1:10 in Triton X-100, and then exposed to two freeze/thaw cycles of 5 minutes in a dry ice/ethanol bath and 2 minutes in 42 °C water bath. Samples are then diluted 1:50 in TE buffer in preparation for use as qPCR template. qPCR is performed using the SYBR Select Master Mix (Applied Biosystems, Waltham, MA) in a Rotor-Gene Q thermocycler (Qiagen, Germantown, MD). PCR primers are specific to the VSVG gene (Forward Primer 5’ GCAAGCATTGGGGAGTCAGAC 3’, Reverse Primer 5’ CTGGCTGCAGCAAAGAGATC 3’). Viral stocks are tested monthly and are typically stable for 12 months when stored at 4 °C and protected from light. While viral genes/ml is a reliable, consistent measurement of viral concentration, the efficiency by which a viral stock successfully transduces mammalian cells varies by cell type, the promoter used to drive expression, and the means by which transduction is detected.

### Molecular Biology

The cDNA for the AT_1_ angiotensin receptor and V_2_ vasopressin receptor were obtained from the cDNA Resource Center (Bloomsburg University, Bloomsburg, PA). The cDNA encoding the β-arrestin2 sensor, Red DAG sensor, R-GECO sensor, Red cADDis sensor, and the receptors were cloned into the same vector which put them under the transcriptional control of a CMV promoter.

### Cell culture and viral transduction

HEK 293T cells were cultured in Eagle’s minimum essential media (EMEM) supplemented with 10% fetal bovine serum (FBS) and penicillin-streptomycin at 37 °C in 5% CO_2_. For BacMam transduction, cells were resuspended in media at a density of 52,000 cells per 100 μL. 100 μL of this suspension was combined with BacMam containing the β-arrestin, Red Downward DAG, Red Upward cADDis, and/or R-GECO sensors and the indicated receptors, 2 mM sodium butyrate, and EMEM in a final volume of 150 μL. For each experiment 4.24 × 10^8^ viral genes of β-arrestin virus were added to each well. For multiplex experiments, 4.79 × 10^8^ viral genes of the Red DAG sensor or 6.18 × 10^8^. Viral genes of the Red cADDis (cAMP) sensor were added with the arrestin sensor. For experiments with the R-GECO sensor, 8.46 × 10^8^ viral genes were added to each well. The viruses carrying the AT_1_ and V_2_ receptors were added such that 2.12 × 10^8^ and 3.04 × 10^8^ viral genes went into each well, respectively. The cell/transduction mixture was then seeded into 96-well plates and incubated for ∼ 24 hours at 37 °C in 5% CO_2_. Thirty minutes prior to fluorescence plate reader or imaging experiments, the media in each well was replaced with 150 μL of Dulbecco’s phosphate buffered saline (DPBS) supplemented with Ca^2+^ (0.9 mM) and Mg^2+^ (0.5 mM).

### Automated plate reader assays

Fluorescence plate reader experiments were performed on the BioTek Synergy Mx (BioTek, Winooski, VT) and BMG CLARIOstar (BMG Labtech, Cary, NC) in 96 well plates. On the Biotek Synergy Mx plate reader, green fluorescence detection was recorded using 488/20 nm excitation and 525/20 nm fluorescence emission, while red fluorescence detection was recorded using 565/20 nm excitation wavelength and 603/20 nm fluorescence emission. On the BMG CLARIOstar plate reader, green fluorescence detection was recorded using 488/14 nm excitation wavelength and 535/30 nm fluorescence emission, while red fluorescence detection was recorded using 566/18 nm excitation wavelength and 620/40 nm fluorescence emission. Drug was added manually with a multichannel pipette in a volume of 50 µL at the indicated time points. While all of the data reported here came from cells in 96 well plates, we and others have been successful using this assay in the 384 well format.

### Drug compounds

Oxytocin and vasopressin were obtained from Cayman Chemical (Ann Arbor, MI). Angiotensin II and SII (Sar^1^, Ile^4,8^]-Angiotensin II) ^39^ were obtained from Genscript (Piscataway, NJ) and MyBioSource (San Diego, CA), respectively. Trevena peptides TRV120026, TRV120045, and TRV120055 ^46^ were synthesized by Genscript. For clarity, the ligand names are abbreviated to TRV026, TRV045 and TRV055. All working concentration of drugs were dissolved in DPBS and added manually to the HEK 293T cells at the indicated concentrations and time points.

### Data analysis

Fluorescence data were normalized to baseline. Specifically, baseline fluorescence i.e. prior to the addition of compound, was measured over at least 5 time points and the average baseline value calculated. The fluorescence value in the well subsequent to the addition of compound or vehicle was divided by the average baseline value for that well, giving the baseline-normalized fluorescence value (Δ F / F). For fitting of the model equations, in which stimulation of signaling was analyzed, downward sensor data were first normalized to be upward (arrestin sensor data in Fig. 2a,b, Fig 4a and Supplementary Fig. S6, and DAG sensor data in Fig. 4b)). This was done by subtracting the baseline-normalized value from unity (1 - ΔF/F). This approach enabled a unified presentation and analysis of stimulation of signaling data.

Curve fitting was performed using Prism 8.1 (Graphpad Software, San Diego, CA). Time course data were fit to exponential equations. The time course data included the baseline phase and the equation incorporated a parameter which represents baseline fluorescence (“y0” or “Baseline” - see below). For downward responses data were fit to the “Plateau followed by one phase decay” equation built into Prism ^57^:

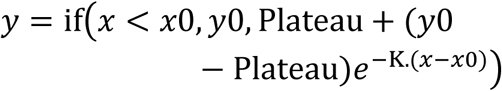

where *y*0 is the baseline signal, *x*0 the time of initiation of the signal, Plateau is the signal at the plateau (formally the asymptote as time approaches infinity) and K the observed rate constant in units of time^-1^. For upward responses data were fit to the “Plateau followed by one phase association” equation built into Prism ^58^:

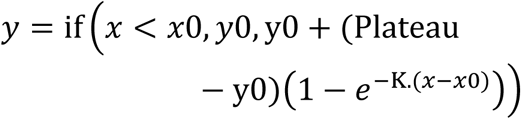

For the rise-and-fall DAG response (Fig. 4b) data were fit to a user-defined bi-exponential equation_24_:

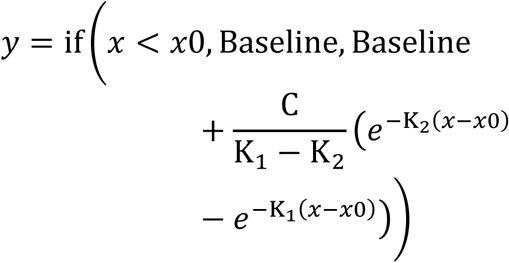

where “Baseline” is the baseline signal (response before addition of ligand), C a fitting constant, and K_1_ and K_2_ rate constants for the two exponential phases in units of time^-1^. This is the general form of the two component signaling model ^24,25^. The calcium response also conformed to a rise-and-fall curve but the baseline response drifted downwards slightly (Fig. 4c). Specifically, the plateau at late time points was slightly lower than the baseline fluorescence prior to the addition of ligand. This drift was incorporated by introducing a drift parameter into the bi-exponential equation:

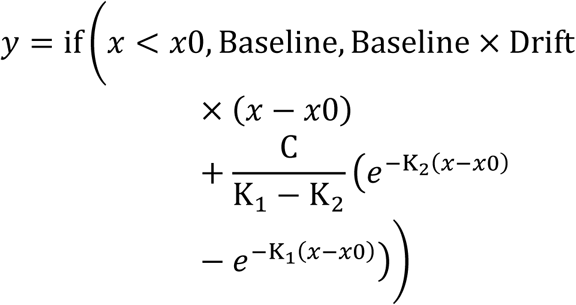

In all the analyses, *x*0, the time of initiation of the signal, was allowed to vary in the analysis (as opposed to being held constant) to accommodate slight differences between wells in the time of addition of ligand.

Concentration-response data were fit to a sigmoid curve equation, the “Log(agonist) vs. response -- Variable slope” equation in Prism ^45^:

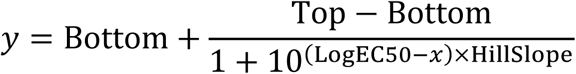

The “Bottom” parameter was constrained to zero in the case of the *k*_*τ*_ analysis, in which the Plateau x *k*_obs_ value is plotted against the agonist concentration.

Technical replicates in the data sets were considered separate points in the curve fitting analysis.

## Appendix Arrestin recruitment mechanism and equations

GPCRs interact with arrestin and this interaction can be detected directly using optical biosensors. In this study the biosensor was a conformationally-sensitive mNeonGreen-tagged arrestin in which the optical properties changed upon binding to the GPCR. The interaction can be described as a straightforward bimolecular interaction between the two proteins, as indicated in the following scheme:

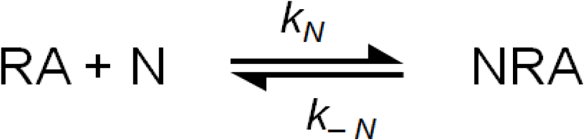

N is arrestin, R is receptor and A is the ligand. RA is receptor-ligand complex and NRA the ternary complex of arrestin, receptor and ligand. *k*_*N*_ is the association rate constant for receptor-arrestin association. The value of this parameter is most likely determined by the rate of receptor phosphorylation since this is the rate-limiting step in receptor-arrestin association. *k*_*-N*_ is the arrestin-receptor dissociation rate constant. *K*_*A*_ is the affinity constant for ligand binding to the receptor (more precisely, the equilibrium dissociation constant). It is assumed that ligand binding is at equilibrium with the receptor, that the unbound receptor (R) does not bind arrestin, and that ligand dissociation from the ternary complex NRA is much slower than that from the binary complex RA.

Equations defining the change of signal (arrestin-receptor complex, NRA) over time were derived. From these equations the initial rate of arrestin recruitment (*k*_*τ*_) emerged as a readily-measurable parameter. Two scenarios regarding stoichiometry were formularized – receptor excess over arrestin (the most likely scenario in this study) and arrestin excess over receptor. Both formulations yield *k*_*τ*_, and *k*_*τ*_ is measured in the same way for both scenarios. Here the equations for arrestin-receptor complex are solved for the two scenarios, then the identity of the initial rate in the equations demonstrated, and finally the method for measuring the initial rate (*k*_*τ*_) presented.

### Receptor excess over arrestin

In this scenario it is assumed receptor is in sufficient excess over arrestin that the arrestin-receptor complex does not appreciably deplete the concentration of receptor. The differential equation defining the change of receptor-arrestin complex over time is,

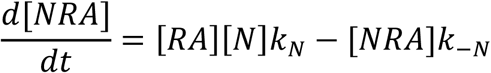

In this case the units of *k*_*N*_ are receptor units^-1^.min^-1^. [RA] can be expressed as a function of the total receptor concentration, as follows. First, since we assume [NRA] does not appreciably deplete the total receptor concentration, the conservation of mass equation for the receptor can be written as,

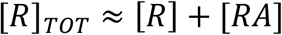

Next, [R] is substituted in this equation. Since ligand-receptor binding is at equilibrium, [R] can be defined as,

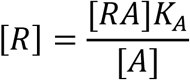

Substituting and rearranging gives the desired expression for [RA]:

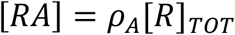

where *ρ*_*A*_ is fractional occupancy of receptor by A, defined by,

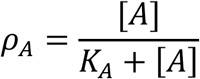

Substituting into the differential equation for [NRA] gives,

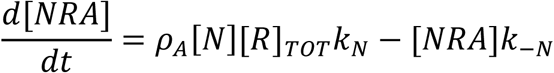

Next the [N] term is substituted with an expression for the total concentration of arrestin. In this scenario, the conservation of mass equation for arrestin is:

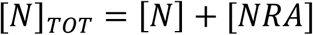

Solving for [N] and substituting into the differential equation gives,

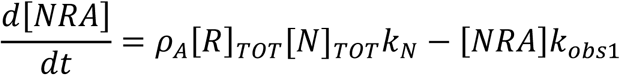

where,

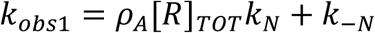

Integrating gives the desired equation defining [NRA] over time, equation (1):

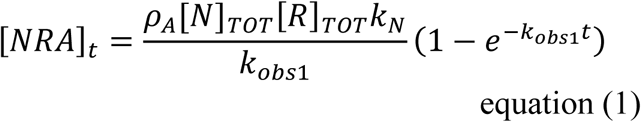

### Arrestin excess over receptor

In this scenario it is assumed arrestin is in sufficient excess over receptor that the arrestin-receptor complex does not appreciably deplete the concentration of arrestin. The differential equation defining the change of receptor-arrestin complex over time is the same as for the receptor excess assumption given above:

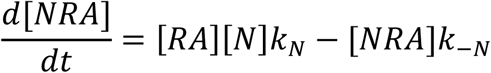

In this case the units of *k*_*N*_ are arrestin units^-1^.min^-1^. Since *N* is in excess over *R*, the free concentration of *N* is approximately equal to the total concentration of *N*, [*N*]_*TOT*_. Consequently, the differential equation can be written as,

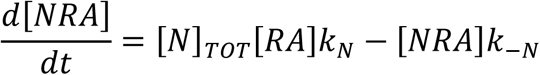

Next, [*RA*] in this equation can be expressed as a function of [*R*]_*TOT*_. The conservation of mass equation is,

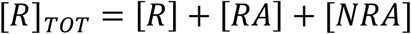

This equation can be rearranged and solved for [RA], utilizing the expression [*R*] = [*RA*]*K*_*A*_/[*A*]

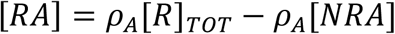

Substituting into the differential equation and rearranging gives,

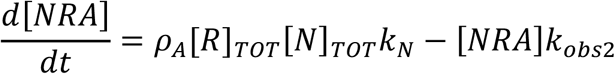

where,

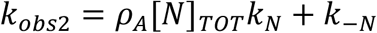

Integrating gives the desired equation defining [NRA] over time, equation (2):

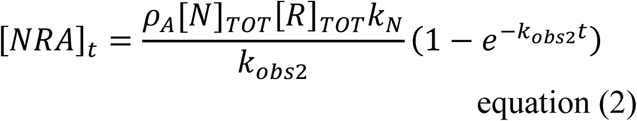

### Defining the initial rate of arrestin recruitment and identifying it in the equations

The initial rate of arrestin interaction with the receptor is that before depletion of arrestin or receptor by formation of the NRA ternary complex, and before breakdown of the complex. This rate is defined as,

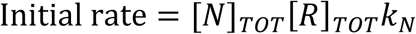

This parameter is a direct analogue of the initial rate of signaling in the kinetic signaling model, the equation for which is,

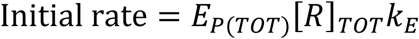

where E_P(TOT)_ is total concentration of precursor and *k*_*E*_ the response generation rate constant. In order to standardize the nomenclature between arrestin recruitment and the signaling models *k*_*τ*_ is used as the term for the initial rate of arrestin recruitment as well as that for signaling:

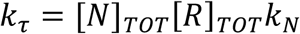

It is evident by visual inspection that this term, [N]_TOT_[R]_TOT_*k*_*N*_ is present in the numerator of the equations for arrestin recruitment over time, equations (1) and (2), reproduced here for convenience:

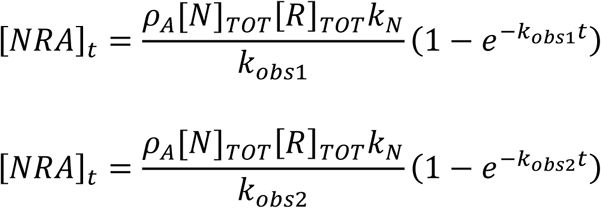

Substituting *k*_*τ*_ for [*N*]_*TOT*_[*R*]_*TOT*_*k*_*N*_ gives the equation used to analyze the time course data, equation (3):

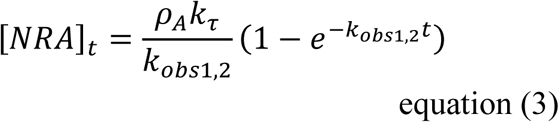

### Estimating the initial rate by curve fitting

Here it is shown that *k*_*τ*_ can be measured by combining parameters from a familiar curve fitting procedure, either from a concentration-response experiment or from an experiment employing a saturating concentration of ligand (Supplementary Fig. S4 and S5, respectively). The equations for both the receptor and arrestin excess assumptions (equations (1) and (2), respectively) take the form of the familiar exponential association equation:

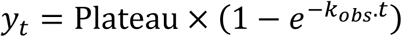

where Plateau is the asymptote, the value of *y* as time approaches infinity, and *k*_*obs*_ the observed rate constant. (This is the “One phase association” equation in GraphPad Prism ^59^). By comparing with equations (1) and (2), it can be seen that the parameters are defined as,

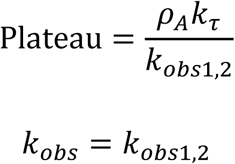

Combining these parameters, by multiplying them together, yields *k*_*τ*_ multiplied by *ρ*_*A*_ (the fractional receptor occupancy by ligand):

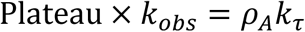

Note that this expression is the same regardless of the excess scenario. It is instructive now to expand the *ρ*_*A*_ term:

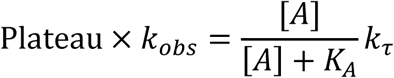

When the ligand concentration is maximally effective for recruiting arrestin, we assume [A] is in large excess over *K*_*A*_. Under this condition, the equation reduces to,

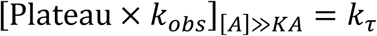

This means that *k*_*τ*_ for arrestin recruitment can be measured by multiplying the plateau by the rate constant for a maximally effective concentration of ligand. This can be done either using a single, maximally stimulating concentration (Supplementary Fig. S5, Fig. 4a) or by plotting the Plateau × *k*_*obs*_ vs [A] (Supplementary Fig. S4, Fig. 2c). In the latter method, the data are analyzed using a sigmoid dose-response equation and in this case *k*_*τ*_ is the fitted maximum Plateau × *k*_*obs*_ value of the curve and *K*_*A*_ is the [A]_50_ concentration, i.e. the value of [A] giving 50% of the maximal Plateau × *k*_*obs*_ value.

## Acknowledgements

Research reported in this publication was supported by National Institute of General Medical Sciences, National Institutes of Health under award number R44GM125390 and National Institute of Neurologic Disorders and Stroke, National Institutes of Health R44NS082222. This content is solely the responsibility of the authors and does not necessarily represent the official views of the National Institutes of Health.

## Author contributions

S.R.J.H. designed the data analysis framework, designed experiments and analyzed data. P.H.T designed and executed experiments and analyzed data. A.M.Q. and T.E.H. designed experiments and conceived the study. All authors wrote the manuscript.

## Supplementary Information

**Supplementary Table 1.**
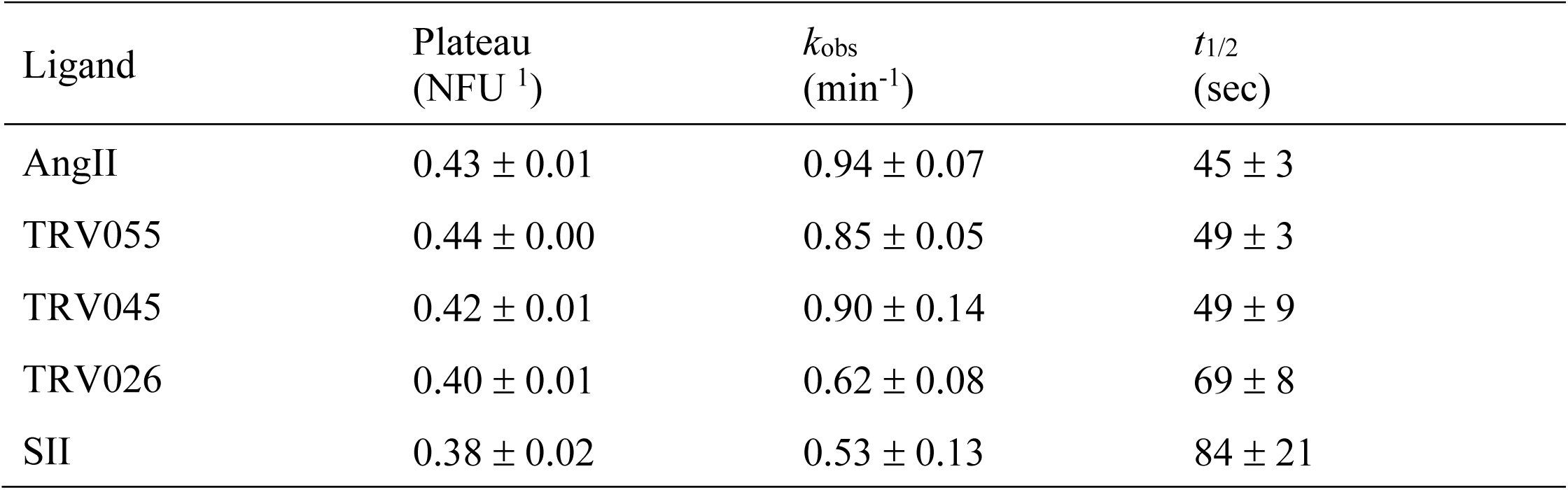
Arrestin recruitment association exponential fit parameter values for the AT1 receptor. Time course data for arrestin recruitment stimulated by the test ligands at 32 μM concentration (see Fig. 2 and Supplementary Fig. S6) was fit to the association exponential equation 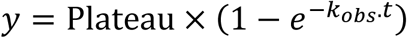 to determine Plateau and *k*_obs_. ^1^ NFU, normalized response units.

**Supplementary Figure S1.**
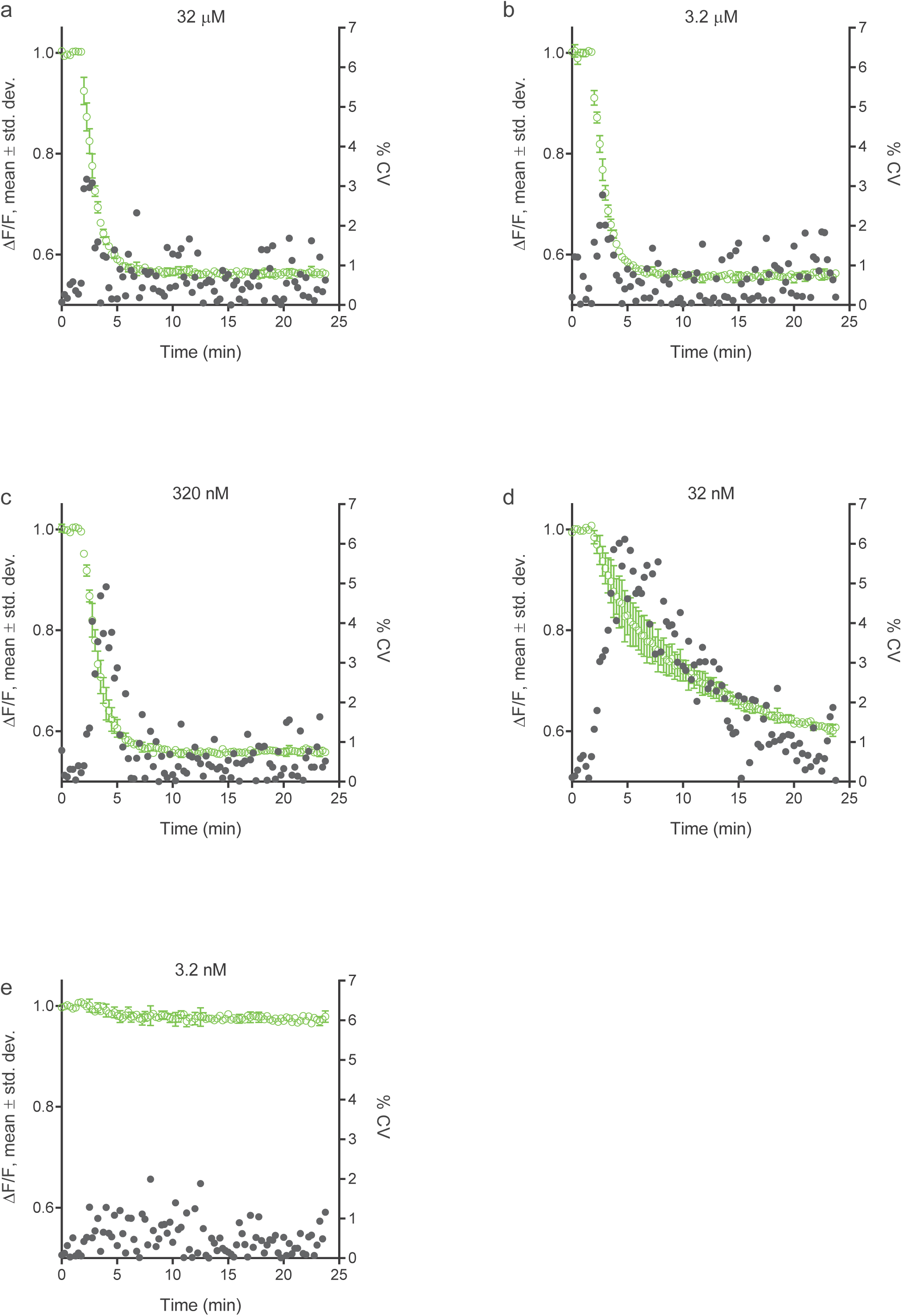
Within-experiment variability between technical replicates, arrestin sensor stimulation by AngII via the AT_1_ receptor. Data points are duplicates, with error bars representing the standard deviation. The % CV was calculated for each data point and is plotted on the right-hand y axis. The analysis was performed for multiple concentrations of AngII, spanning the effective range (a, 32 μM; b, 3.2 μM; c, 320 nM; d, 32 nM; e, 3.2 nM). The time interval between data points was 15 seconds. Fluorescence intensity was measured using the BMG BMG CARIOstar and data normalized to baseline fluorescence before agonist addition.

**Supplementary Figure S2.**
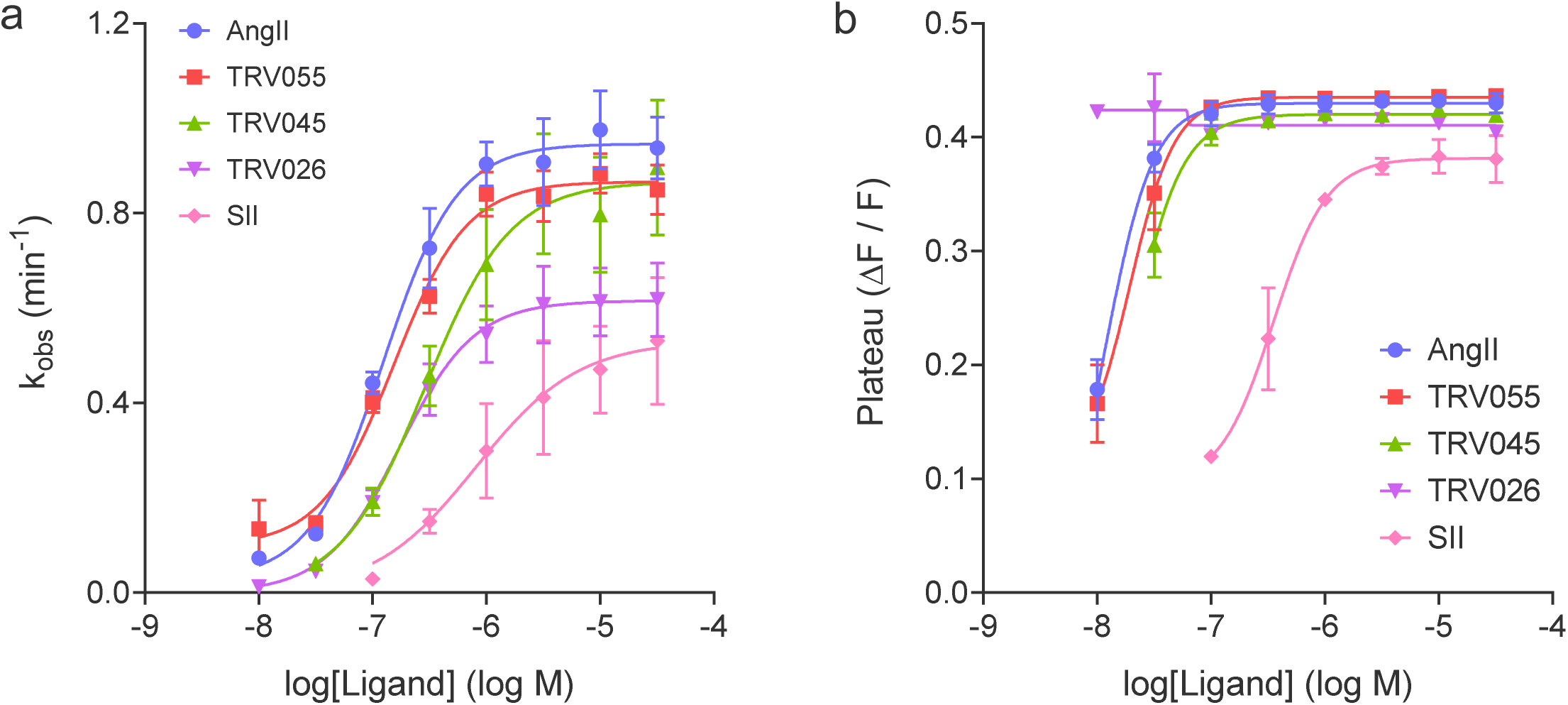
Curve fit parameters for the time course of arrestin recruitment to the AT_1_ angiotensin receptor by various ligands: (a) *k*_*obs*_, (b) Plateau. The time course data (Fig. 2a and b, Supplementary Fig. S6) were fit to the association exponential equation:

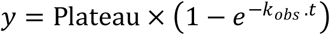 The measured values of the observed rate constant *k*_obs_ and Plateau are plotted against the ligand concentration. Note the value of both parameters increases as the ligand concentration increases. The curves are fits to a sigmoid dose-response equation.

**Supplementary Figure S3.**
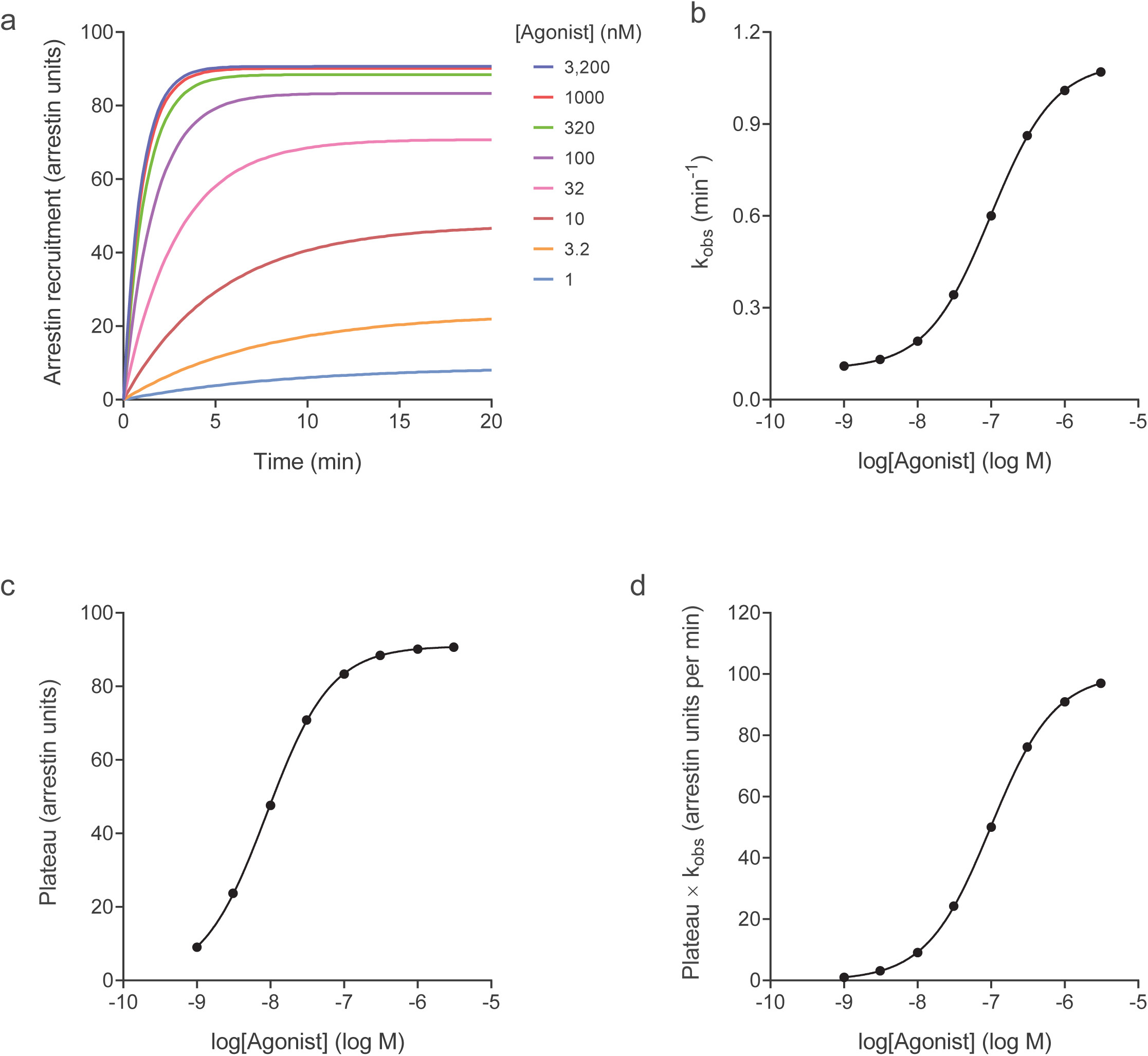
Simulated arrestin recruitment data. The arrestin recruitment model (Appendix) was used to simulate the time course of arrestin recruitment to a GPCR stimulated by agonist ligand. Time course data (a) were simulated using equation (1), assuming receptor is in excess over arrestin, for a range of concentrations of agonist. From these data the observed rate constant *k*_*obs*_ and Plateau can be calculated using equation (3). The calculated values are plotted versus the agonist concentration (b, *k*_*obs*_; c, Plateau). Note the value of both parameters increases as the ligand concentration increases. The curves are fits to a sigmoid dose-response equation. Finally, the initial rate of recruitment at each concentration can be determined by multiplying the Plateau by *k*_*obs*_. The resulting value is plotted against the agonist concentration (d). The data are then fit by the sigmoid dose-response equation to determine *k*_*τ*_ (*k*_*obs*_ × Plateau at maximally-effective [A]) and agonist affinity for the receptor ([A]_50_). Model parameter values used for the simulation were: *K*_*A*_, 100 nM; [R]_TOT_, 100 receptor units; [N]_TOT_, 100 arrestin units; *k*_*N*_, 0.01 receptor units^-1^min^-1^; *k*_*-N*_, 0.1 min^-1^. The calculated *k*_*τ*_ value was 100 arrestin units.min^-1^.

**Supplementary Figure S4.**
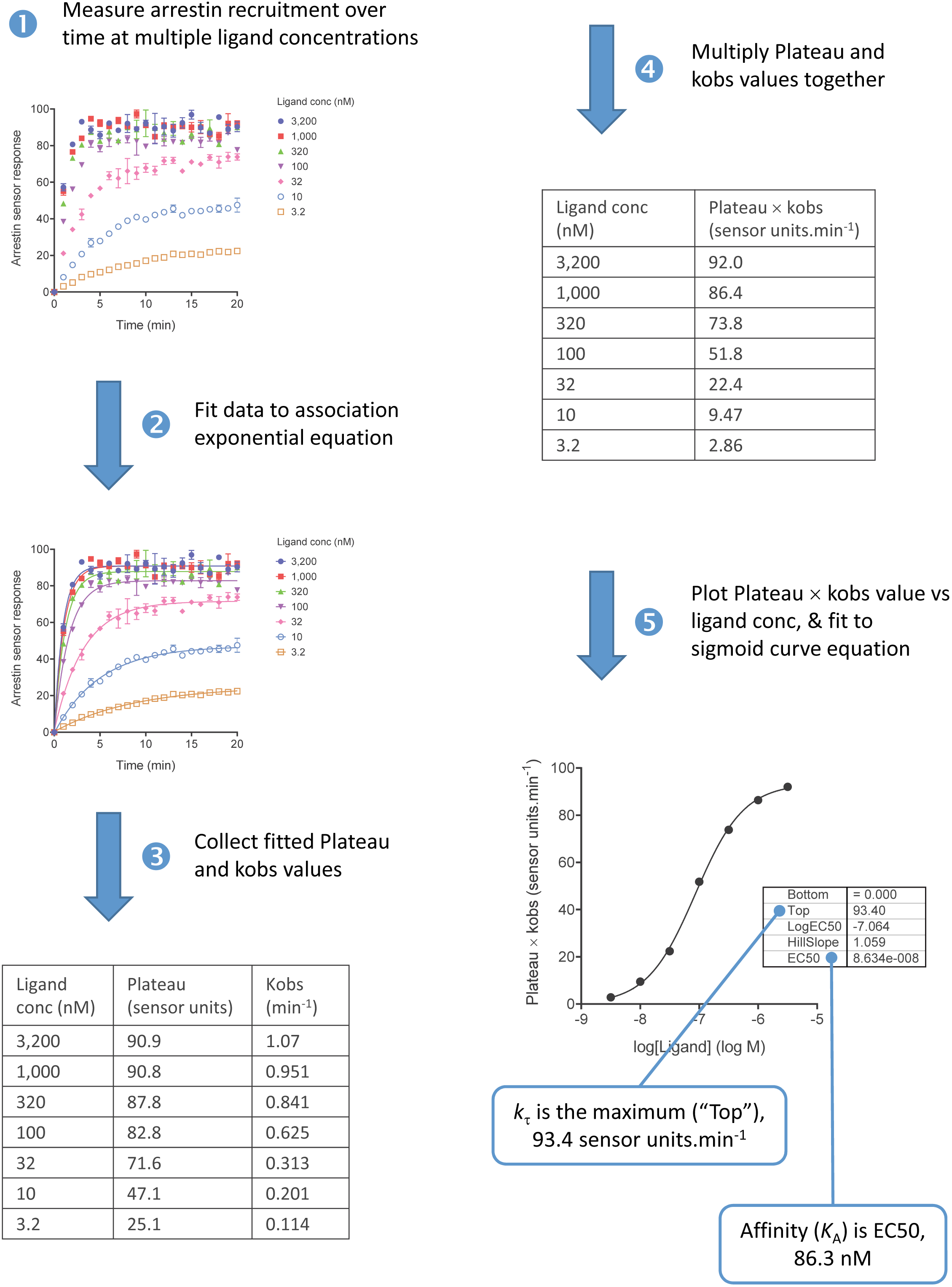
Quantifying *k*_*τ*_ concentration response method

**Supplementary Figure S5.**
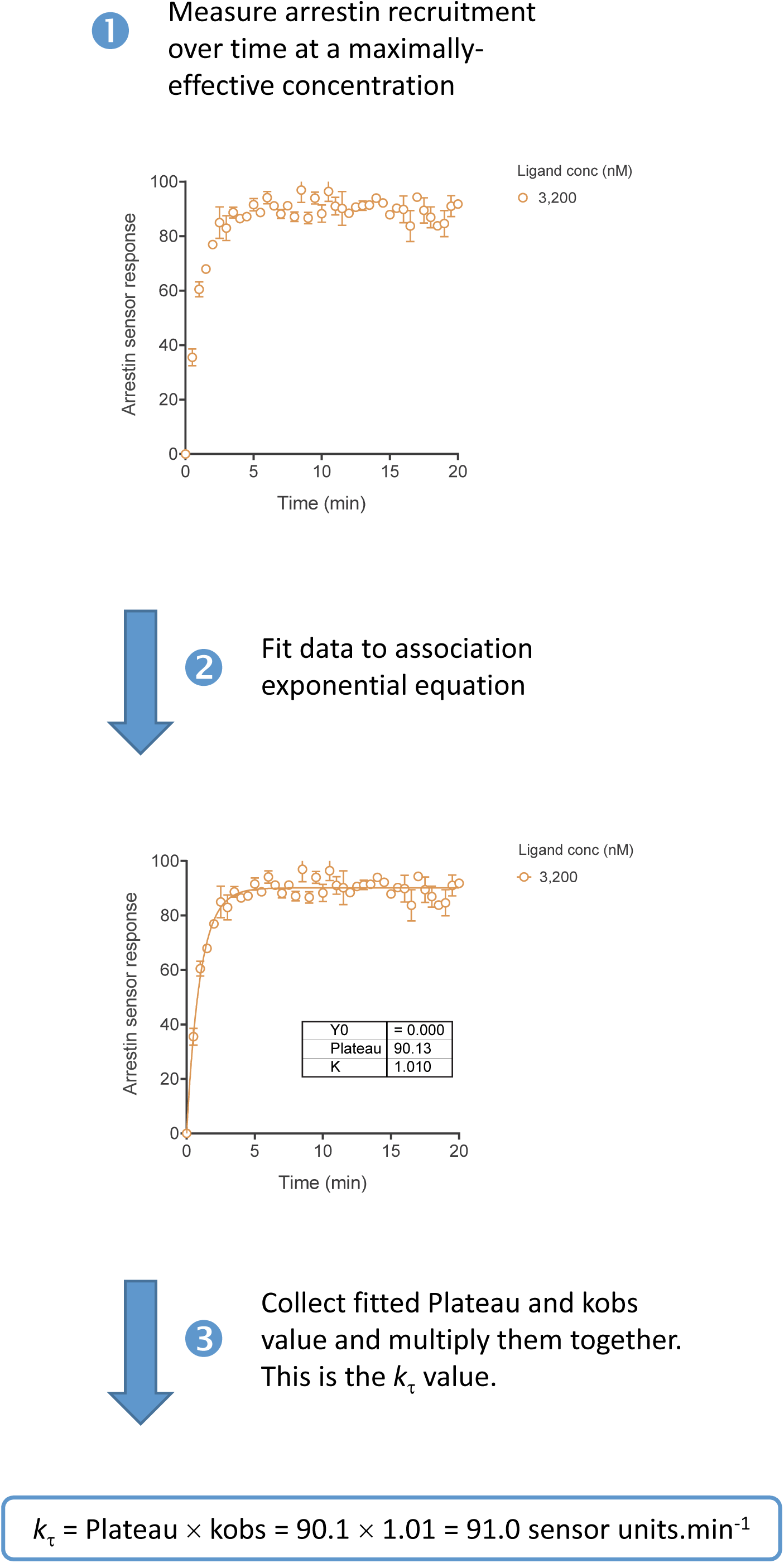
Quantifying *k*_*τ*_ maximally-stimulating concentration method

**Supplementary Figure S6.**
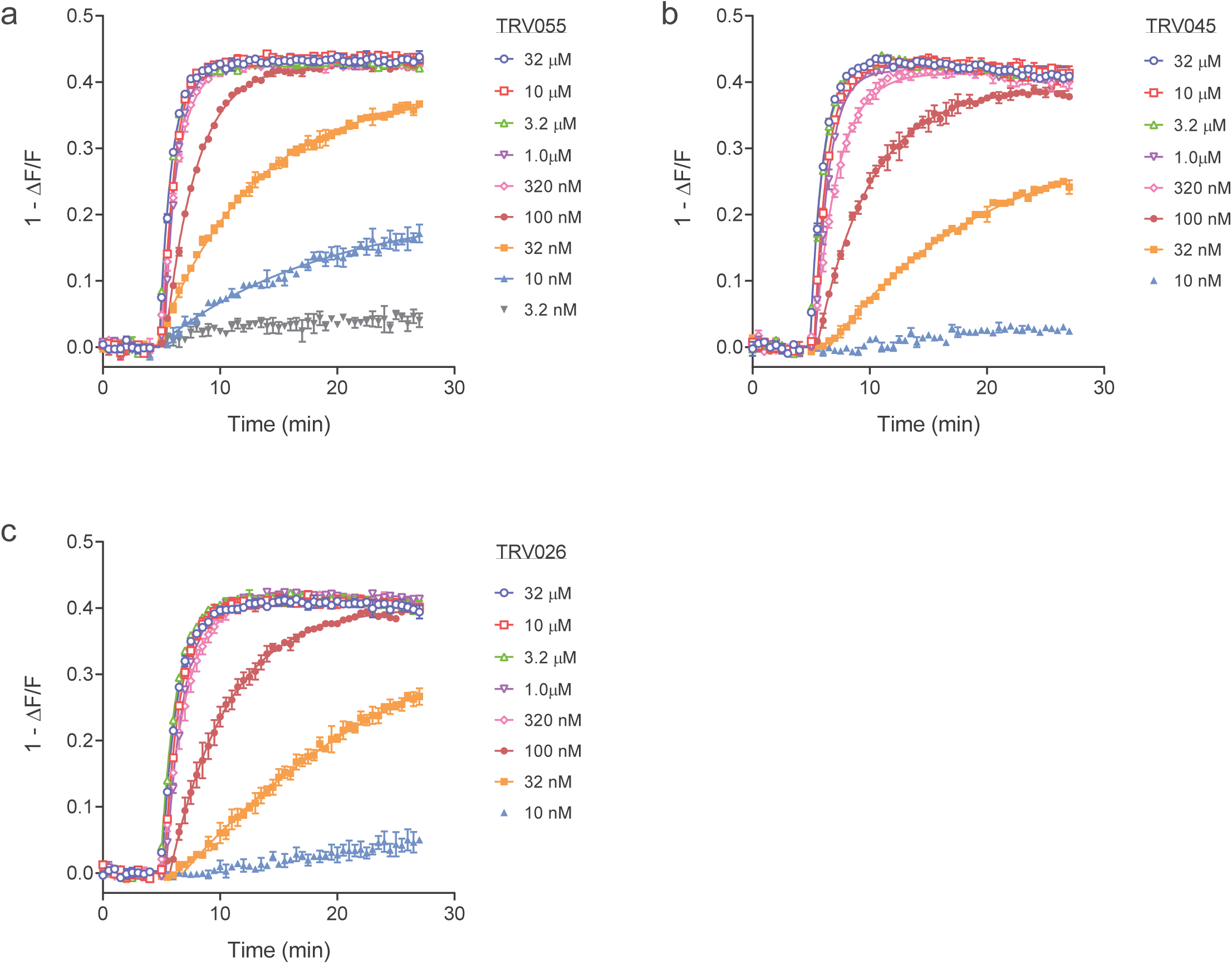
Time course concentration response of arrestin recruitment to the AT_1_ angiotensin receptor stimulated by TRV055 (a), TRV045 (b) and TRV026 (c). Curves are the fits to the association exponential equation:

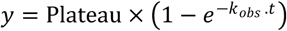 From the fitted value of *k*_*obs*_ and Plateau *k*_*τ*_ can be calculated as described in Fig. 2. Data are from the Biotek Synergy Mx plate reader.

